# Global Root Traits (GRooT) Database

**DOI:** 10.1101/2020.05.17.095851

**Authors:** Nathaly Guerrero-Ramirez, Liesje Mommer, Grégoire T. Freschet, Colleen M. Iversen, M. Luke McCormack, Jens Kattge, Hendrik Poorter, Fons van der Plas, Joana Bergmann, Thom W. Kuyper, Larry M. York, Helge Bruelheide, Daniel C. Laughlin, Ina C. Meier, Catherine Roumet, Marina Semchenko, Christopher J. Sweeney, Jasper van Ruijven, Oscar J. Valverde-Barrantes, Isabelle Aubin, Jane A. Catford, Peter Manning, Adam Martin, Rubén Milla, Vanessa Minden, Juli G. Pausas, Stuart W. Smith, Nadejda A. Soudzilovskaia, Christian Ammer, Bradley Butterfield, Joseph Craine, Johannes H.C. Cornelissen, Franciska T. de Vries, Marney E. Isaac, Koen Kramer, Christian König, Eric G. Lamb, Vladimir G. Onipchenko, Josep Peñuelas, Peter B. Reich, Matthias C. Rillig, Lawren Sack, Bill Shipley, Leho Tedersoo, Fernando Valladares, Peter van Bodegom, Patrick Weigelt, Justin P. Wright, Alexandra Weigelt

## Abstract

**Motivation:** Trait data are fundamental to quantitatively describe plant form and function. Although root traits capture key dimensions related to plant responses to changing environmental conditions and effects on ecosystem processes, they have rarely been included in large-scale comparative studies and global models. For instance, root traits remain absent from nearly all studies that define the global spectrum of plant form and function. Thus, to overcome conceptual and methodological roadblocks preventing a widespread integration of root trait data into large-scale analyses we created the Global Root Trait (GRooT) Database. GRooT provides ready-to-use data by combining the expertise of root ecologists with data mobilization and curation. Specifically, we (i) determined a set of core root traits relevant to the description of plant form and function based on an assessment by experts, (ii) maximized species coverage through data standardization within and among traits, and (iii) implemented data quality checks.

**Main types of variables contained:** GRooT contains 114,222 trait records on 38 continuous root traits.

**Spatial location and grain:** Global coverage with data from arid, continental, polar, temperate, and tropical biomes. Data on root traits derived from experimental studies and field studies.

**Time period and grain:** Data recorded between 1911 and 2019

**Major taxa and level of measurement:** GRooT includes root trait data for which taxonomic information is available. Trait records vary in their taxonomic resolution, with sub-species or varieties being the highest and genera the lowest taxonomic resolution available. It contains information for 184 sub-species or varieties, 6,214 species, 1,967 genera and 254 families. Due to variation in data sources, trait records in the database include both individual observations and mean values.

**Software format:** GRooT includes two csv file. A GitHub repository contains the csv files and a script in R to query the database.

## 1. Introduction

Plant traits have been used for describing multiple aspects of plant species’ fitness and realized performance, including growth, survival, and reproduction (Grime, 1977; Calow, 1987; Geber & Griffen, 2003; Reich *et al*., 2003; Adler *et al*., 2014; Díaz *et al*., 2016). Moreover, traits can illustrate how species respond to environmental variability and disturbances (Grime, 1974; Keddy, 1992; Pausas *et al*., 2004; Bruelheide *et al*., 2018; Minden & Olde Venterink, 2019; Wieczynski *et al*., 2019) and reveal species effects on ecosystem functions (Díaz & Cabido, 2001; Lavorel & Garnier, 2002; Breitschwerdt *et al*., 2018; Craven *et al*., 2018). While root traits are likely to capture key dimensions of plant form and function, plant evolutionary history, and responses to environmental variability (Bardgett *et al*., 2014; Laliberté, 2016; Freschet *et al*., 2017; Valverde-Barrantes *et al*., 2017; Ma *et al*., 2018; Kong *et al*., 2019), they remain underrepresented in large-scale comparative studies and global models. Accordingly, root traits remain absent from nearly all existing studies that define the global spectrum of plant form and function (Wright *et al*., 2004; Chave *et al*., 2009; Reich, 2014; Díaz *et al*., 2016) but see Averill *et al*., (2019).

Conceptual and methodological challenges have deterred widespread data integration of root traits into global trait databases. Conceptually, the functional importance of some root traits has yet to be formally established, which may preclude their use in large-scale analyses (Aubin *et al*., 2016). Methodologically, quantifying root traits is labor-intensive and there are technical difficulties in obtaining accurate measurements (e.g., Delory *et al*., 2017). Further, large variation in methodologies precludes data standardization and integration within traits. Specifically, while traits are characteristics measurable at the individual plant level (Violle *et al*., 2007); root traits may be measured in different ways, increasing the number of trait variables. For example, data for root nitrogen uptake are separated into eight trait variables (Iversen *et al*., 2017). While coordinated initiatives such as the Fine-Root Ecology Database (FRED; Iversen *et al*., 2017) and the Plant Trait Database (TRY; Kattge *et al*., 2011, 2020) have compiled valuable root trait data, these databases still face many of these conceptual and methodological challenges associated with root traits.

FRED has been essential in terms of fine-root trait data mobilization, being the largest contributor of root trait data to TRY (Kattge *et al*., 2020). FRED contains approximately 300 root trait variables; this high resolution of root variables allows users to investigate a broad set of research questions. However, barriers remain when using these root trait data in the context of large-scale comparative studies. For example, the number of trait variables can be overwhelming, particularly for non-root specialists. Further, a large number of trait variables have few data records, limiting data quality checks. For example, TRY performs data standardization and intensive data quality checks for traits with more than 1,000 records (Kattge *et al*., 2020), yet most root traits have fewer records than this threshold. In addition, some trait variables that are not directly comparable in terms of definitions and units have been aggregated by type on TRY, for example, root type/root architecture. Therefore, using these data requires that one first disaggregates these traits (e.g., by establishing links between trait names and definitions) and then standardizes trait values. Finally, accurate global assessments on root trait data availability, in terms of geographic or phylogenetic coverage, are essential to identify data gaps and to work towards increasing representativeness in large-scale comparative studies and dynamic global vegetation models.

To overcome these roadblocks, we have created the Global Root Trait (GRooT) Database. The main objective of GRooT is to make root trait data ready-to-use, particularly in the context of large-scale analyses. To do so, we first provide a set of core root traits that are considered to be relevant for describing plant form and function. Trait selection builds on the compilation of standardized trait measurements in a new handbook on root traits (Freschet *et al*., submitted) and an assessment by experts on root traits. In addition, we improved data coverage by compiling information from existing databases, mobilizing new data and standardizing data across methodologies within and among traits. Further, we curated and performed data quality checks for each root trait in GRooT and make these data publicly available. Secondly, we provide within GRooT a unique overview of global root trait availability in terms of geographic and phylogenetic coverage. We envision that our advanced root trait database will be informative to global trait-based models and help to guide future measurement initiatives.

## 2. Methods

### 2.1 Data acquisition and compilation

GRooT includes root trait data provided directly from researchers, extracted from literature, or from large databases such as FRED version 2.3 (https://roots.ornl.gov/; Iversen *et al*., 2017, 2018) and TRY version 4.1 (https://www.try-db.org; Kattge *et al*., 2011, 2020). In total, GRooT includes data from 919 publications via FRED, 38 datasets via TRY and 12 additional datasets (Appendix 1, Data References).

GRooT was assembled by first determining which root traits are most relevant in terms of describing plant form and function (Table 1 and Supporting information Tables S1 and S2). To build towards an ontology of root traits, we standardized trait names across data sources (Supporting information Table S3) and matched them with names from the new handbook of root traits (Freschet *et al*., submitted).

**Table 1.** Root traits included in GRooT. For each trait, standardized units, number of species, and number of species-by-site mean values are presented. Traits are categorized based on McCormack et al. (2017) and Freschet et al. (submitted). See Supporting information Table S1 for trait definitions.

| Traits | Units | # species | # Species by site | Mean | Qu. (0.25) | Median | Qu. (0.75) |
| --- | --- | --- | --- | --- | --- | --- | --- |
| **Anatomy** |  |  |  |  |  |  |  |
| Root cortex thickness | μm | 151 | 180 | 207.8 | 76.9 | 153.2 | 300.4 |
| Root stele diameter | μm | 318 | 491 | 185.9 | 57.9 | 104.1 | 204.4 |
| Root stele fraction | % | 352 | 534 | 0.13 | 0.05 | 0.08 | 0.15 |
| Root xylem vessel number<sup>t</sup> | number mm<sup>-2</sup> | 96 | 97 | 4438 | 2325 | 3828 | 5847 |
| Root vessel diameter<sup>t</sup> | μm | 125 | 125 | 9.2 | 4.2 | 6.5 | 9.1 |
| **Architecture** |  |  |  |  |  |  |  |
| Root branching ratio | number number<sup>-1</sup> | 173 | 189 | 3.52 | 2.45 | 3.12 | 4.30 |
| Root branching density<sup>‡</sup> | number cm<sup>-1</sup> | 216 | 247 | 2.79 | 1.62 | 2.17 | 3.35 |
| **Belowground allocation** |  |  |  |  |  |  |  |
| Root mass fraction | g g<sup>-1</sup> | 1348 | 3527 | 0.40 | 0.24 | 0.36 | 0.54 |
| **Chemistry** |  |  |  |  |  |  |  |
| Root structural carbohydrate concentration | mg g<sup>-1</sup> | 185 | 228 | 567.9 | 443.1 | 620.0 | 692.6 |
| Root lignin concentration | mg g<sup>-1</sup> | 311 | 401 | 164.2 | 96.5 | 153.5 | 203.2 |
| Root carbon concentration | mg g<sup>-1</sup> | 1099 | 2328 | 417.3 | 388.2 | 416.7 | 455.5 |
| Root nitrogen concentration | mg g<sup>-1</sup> | 1719 | 3619 | 12.5 | 7.8 | 10.8 | 15.5 |
| Root phosphorus concentration | mg g<sup>-1</sup> | 486 | 1284 | 1.33 | 0.74 | 1.10 | 1.61 |
| Root carbon-to-nitrogen ratio | mg mg<sup>-1</sup> | 925 | 1308 | 41.6 | 25.3 | 35.8 | 51.5 |
| Root nitrogen-to-phosphorus ratio | mg mg<sup>-1</sup> | 154 | 154 | 13.7 | 6.0 | 10.6 | 19.0 |
| Root calcium concentration | mg g<sup>-1</sup> | 169 | 915 | 5.10 | 2.44 | 3.38 | 5.57 |
| Root potassium concentration | mg g<sup>-1</sup> | 167 | 891 | 6.88 | 3.65 | 5.95 | 8.92 |
| Root magnesium concentration | mg g<sup>-1</sup> | 146 | 862 | 2.55 | 1.39 | 2.09 | 3.27 |
| Root manganese concentration | mg g<sup>-1</sup> | 52 | 89 | 0.22 | 0.10 | 0.16 | 0.29 |
| **Dynamics & decomposition** |  |  |  |  |  |  |  |
| Root production<sup>*</sup> | g m<sup>-2</sup> yr<sup>-1</sup> | 116 | 201 | 397.5 | 116.6 | 230.0 | 436.8 |
| Root lifespan mean | days | 80 | 98 | 696.2 | 252.5 | 377.0 | 734.5 |
| Root lifespan median | days | 60 | 65 | 324.8 | 149.5 | 255.7 | 357.9 |
| Root turnover rate | yr<sup>-1</sup> | 126 | 251 | 0.79 | 0.21 | 0.55 | 1.04 |
| Root litter mass loss rate | yr<sup>-1</sup> | 232 | 289 | 1.92 | 0.27 | 0.67 | 2.51 |
| **Horizontal plant mobility** |  |  |  |  |  |  |  |
| Lateral spread<sup>**</sup> | cm yr<sup>-1</sup> | 1398 | 1398 | 7.92 | 0.50 | 6.80 | 13.0 |
| **Microbial associations<sup>\*</sup>** |  |  |  |  |  |  |  |
| Root mycorrhizal colonization intensity<sup>**</sup> | % | 2405 | 2529 | 40.5 | 9.4 | 40.0 | 64.5 |
| **Morphology** |  |  |  |  |  |  |  |
| Mean root diameter | mm | 1628 | 2845 | 0.44 | 0.22 | 0.36 | 0.55 |
| Root dry matter content | g g<sup>-1</sup> | 431 | 1499 | 0.22 | 0.15 | 0.22 | 0.28 |
| Root tissue density | g cm<sup>-3</sup> | 1465 | 3291 | 0.25 | 0.11 | 0.20 | 0.34 |
| Specific root area | cm<sup>2</sup> g<sup>-1</sup> | 477 | 707 | 617.7 | 179.1 | 385.8 | 728.4 |
| Specific root length | m g<sup>-1</sup> | 1973 | 3407 | 88.7 | 18.9 | 47.4 | 113.0 |
| ***Physiology & respiration*** |  |  |  |  |  |  |  |
| Net uptake rate of nitrogen | μmol g<sup>-1</sup>day<sup>-1</sup> | 68 | 68 | 49.5 | 32.3 | 52.6 | 64.8 |
| Specific root respiration | nmol g<sup>-1</sup>s<sup>-1</sup> | 248 | 408 | 21.7 | 7.0 | 14.6 | 32.7 |
| ***System & distribution*** |  |  |  |  |  |  |  |
| Fine root mass-to-leaf mass ratio | g g<sup>-1</sup> | 143 | 176 | 2.39 | 0.08 | 0.32 | 2.30 |
| Coarse-to-fine root mass ratio | g g<sup>-1</sup> | 128 | 130 | 10.6 | 1.7 | 5.0 | 12.8 |
| Root mass density | g cm<sup>-3</sup> | 152 | 260 | 0.13 | 0.0001 | 0.02 | 0.13 |
| Root length density | cm cm<sup>-3</sup> | 122 | 178 | 4.10 | 0.43 | 1.82 | 5.80 |
| Maximum rooting depth | m | 1024 | 1782 | 2.21 | 0.50 | 1.10 | 2.00 |

<sup>†</sup>This information can be used to calculate theoretical root specific hydraulic conductance (Valenzuela-Estrada *et al.*, 2009).
<sup>‡</sup>This information needs to be interpreted with caution, since the included total root length can vary across studies.
<sup>\*</sup>This trait can be measured via minirhizotrons or ingrowth cores and both measurements lead to contrasting results.
<sup>\*\*</sup>Lateral spread by clonal growth: while this trait is not categorized as a trait of the root system *per se*, was included because its influence on root growth (Klimešová & Bello, 2009).
<sup>\*</sup>Qualitative microbial association traits including mycorrhizal association type and nitrogen fixing capacity are included in GRooT (see Supporting information Table S5).
<sup>\*\*</sup>Mycorrhizal colonization intensity is based mostly on data for arbuscular mycorrhizal colonization.

Subsequently, we checked available trait variables (> 700) to establish (i) which variable was associated with pre-selected relevant root traits, (ii) which variables would be the most pertinent for each root trait based mostly on the handbook of root traits; (Freschet *et al*., submitted), and (iii) which variables could be standardized across methodologies within or among traits. Within traits, we aggregated comparable trait variables into a single unique trait, e.g., specific root respiration was combined into a unique trait, independent of it being measured as O_2_ consumption or CO_2_ release (Supporting information Table S3). Among traits, we re-calculated values for traits that could be standardized, such as (i) data on the root-to-shoot ratio for the calculation of root mass fraction (RMF) and (ii) data on stele diameter for the calculation of the stele fraction (Supporting information Table S3 and Fig S1). After this process, we retained those relevant traits with data for more than 50 plant species in the database (Table 1), because traits with lower species coverage seemed less helpful for large-scale analyses involving many species. Traits under this threshold – while still relevant – are currently excluded from GRooT (Supporting information Table S2).

In GRooT, we only included trait records for which taxonomic information was available and excluded trait records where data was taken at the community level, i.e., from species mixtures. Trait records varied in their taxonomic resolution, with sub-species or varieties and genera being the highest and lowest taxonomic resolution available, respectively. We used the generic term of “root”, which includes any kind of root entity (e.g. established using either diameter cut-offs, orders, or functionality). While the need to analyse root entities separately (e.g., separating between fine and coarse roots; root orders or diameter cut-offs; or absorptive and transport roots) is generally recommended by a range of recent syntheses (McCormack *et al*., 2015; Freschet & Roumet, 2017), which entity is most suitable may vary strongly depending on the research question (Freschet & Roumet, 2017). Therefore, we have included information in GRooT that allows one to select data based on root entities (Supplementary information Table S4). We urge future data contributors to provide information about root entities and data users to consider this issue carefully.

GRooT includes selected meta-data for each trait record, when available, such as taxonomic information, experimental conditions, sampling procedure, geographic location, date, as well as climatic and soil variables (Supporting information Table S5). Moreover, we have included additional information for each trait record including species growth form, photosynthetic pathway, and woodiness (Supporting information Table S5). We extracted this information from TRY and the Global Inventory of Floras and Traits (GIFT; Weigelt *et al*., 2019), or from general web research (e.g., Flora of China (www.efloras.org), SEINet (swbiodiversity.org), USDA (plants.usdy.gov, and Southwest Desert Flora (southwestdesertflora.com)) when the information was not available in the aforementioned databases. We also included the present or absent ability to grow clonally and bud bearing information at the species level on GRooT based on the CLO-PLA Database (CLO-PLA; http://clopla.butbn.cas.cz/; Klimešová & Bello, 2009; Klimešová *et al*., 2017, 2019). For data collected under field conditions, biome classification according to Köppen-Geiger was included using the “kgc” R Package (Bryant *et al*., 2017).

We added information on qualitative root traits as mycorrhizal association type and nitrogen (N_2_)-fixing capacity by interconnecting existing databases. For mycorrhizal type, we extracted data from the “FungalRoot: Global online database of plant mycorrhizal associations” (Soudzilovskaia *et al*.). Mycorrhizal assignments were made at the genus level for plant species for which the mycorrhizal status is, according to current knowledge, conserved at this level (Soudzilovskaia *et al*.). We included both, standardized mycorrhizal types (named: mycorrhizalAssociationTypeFungalRoot) and mycorrhizal type from the original source (named: mycorrhizalAssociationType) in the database. For N_2_-fixation capacity, we extracted data from the “Global database of plants with root-symbiotic nitrogen fixation: NodDB Database” (version 1.3a; Tedersoo *et al*., 2018) at the genus level.

### 2.2. Data curation and quality control

We cross-checked references associated with each dataset to avoid data redundancy, which was mostly generated by (i) a dataset being submitted to multiple databases or (ii) databases including the dataset from different sources, e.g., submitted by the main authors versus extracted from literature, or the same data being used in multiple papers. When datasets or references appeared in multiple data sources, we performed manual checks to ensure the removal of redundant measurements while ensuring that complementary information was not removed. In some cases, data contributors were contacted directly to avoid dataset overlaps. Despite these efforts, there is the possibility that some redundant information remains in GRooT, which is most likely restricted to instances where data has been used in multiple publications.

GRooT contains original species names (as provided by the main source) and standardized species names. We standardized original species names using the Taxonomic Name Resolution Service version 4.0, i.e., TNRS, (http://tnrs.iplantcollaborative.org/; Accessed: September 2019; Boyle *et al*., 2013), selecting the best match among The Plant List version 1.1. (http://www.theplantlist.org/; accessed: 19 Aug 2015), Global Compositae Checklist (GCC; http://compositae.landcareresearch.co.nz/Default.aspx; accessed: 21 Aug 2015), International Legume Database and Information Service(ILDIS; http://www.ildis.org/; accessed: 21 Aug 2015), Missouri Botanical Garden’s Tropicos Database (http://www.tropicos.org, accessed 19 Dec 2014), and the USDA’s Plant Database (http://plants.usda.gov, accessed: 17 Jan 2015). In addition, we obtained plant taxonomic order from “taxize” R package version 0.9.4 (Chamberlain & Szöcs, 2013).

We checked trait records to ensure that data were in standardized units. Potential mistakes were checked in the original sources, corrected when possible or excluded when values were unreasonable (e.g., negative values for nutrient concentrations or percentages greater than 100). We calculated the error risk as the number of mean standard deviations (across all species within trait) from the respective species mean (named: errorRisk), following TRY protocol (TRY; Kattge *et al*., 2011, 2020) – but not implemented across root traits in TRY. We reported the number of data entries used to calculate the error risk per species (named: ErrorRiskEntries), with error risk robustness increasing when based on multiple replicates (preferable more than 10 data entries). Normal distribution was checked for each trait and logarithmic transformations were used before calculating error risk scores when required. Large error risk scores can indicate potential measurement errors. Yet they can also reflect intra-specific variation. Thus, we did not use error risk scores to remove trait records from the database but provide them to be used at the users’ discretion.

### 2.3 Data use guidelines and data availability

GRooT contains two csv files and an R script. The first csv file named GRooTFullVersion.csv provides root trait data at the highest resolution available (either trait values from individual replicates or mean values per study), information to filter data by entities (Supporting information Table S4), meta-data (Supporting information Table S5), and error risk scores. The second file named GRooTAggregateSpeciesVersion.csv provides mean, median, and quantiles (0.25 and 0.75) of species values. The R script named GRooTExtraction includes code to calculate error risk and the steps to calculate the mean, median and quantiles (0.25 and 0.75) of species values. The code of the R script is customizable, including options to calculate mean values by excluding trait records based on the error risk, and to select data based on root entities (Supporting information Table S4) or relevant covariables such as root vitality (McCormack *et al*., 2015). GRooT is public and will be maintained in a GitHub repository (https://github.com/GRooT-Database/GRooT-Data). We encourage users to report mistakes, suggestions to improve the database and contribute data (https://groot-database.github.io/GRooT/).

## 3. Description of the data

GRooT includes 38 root traits, 38,276 species-by-site mean values based on 114,222 trait records (Table 1, Supporting information Figs S2 to S14). GRooT includes more than 1,000 species with data on the following nine traits: root mass fraction, root carbon and nitrogen concentration, lateral spread, root mycorrhizal colonization intensity, mean root diameter, root tissue density, specific root length, and maximum rooting depth. Data were collected in experimental microcosm studies (20.2% and 1.3% of species-by-site mean values from potted and hydroponic experiments, respectively), field studies (71.4% of species-by-site mean values; including field observations and field or common garden experiments), or were unspecified (7.1% of species-by-site mean values). Root trait coverage from field studies varies geographically across the globe (Fig 1). Regions such as North America, Europe, and Asia are well-covered, while there are consistent gaps in other regions such as Africa and South America. These geographic patterns are observed in terms of number of species as well as number of traits measured per site.

**Figure 1.**
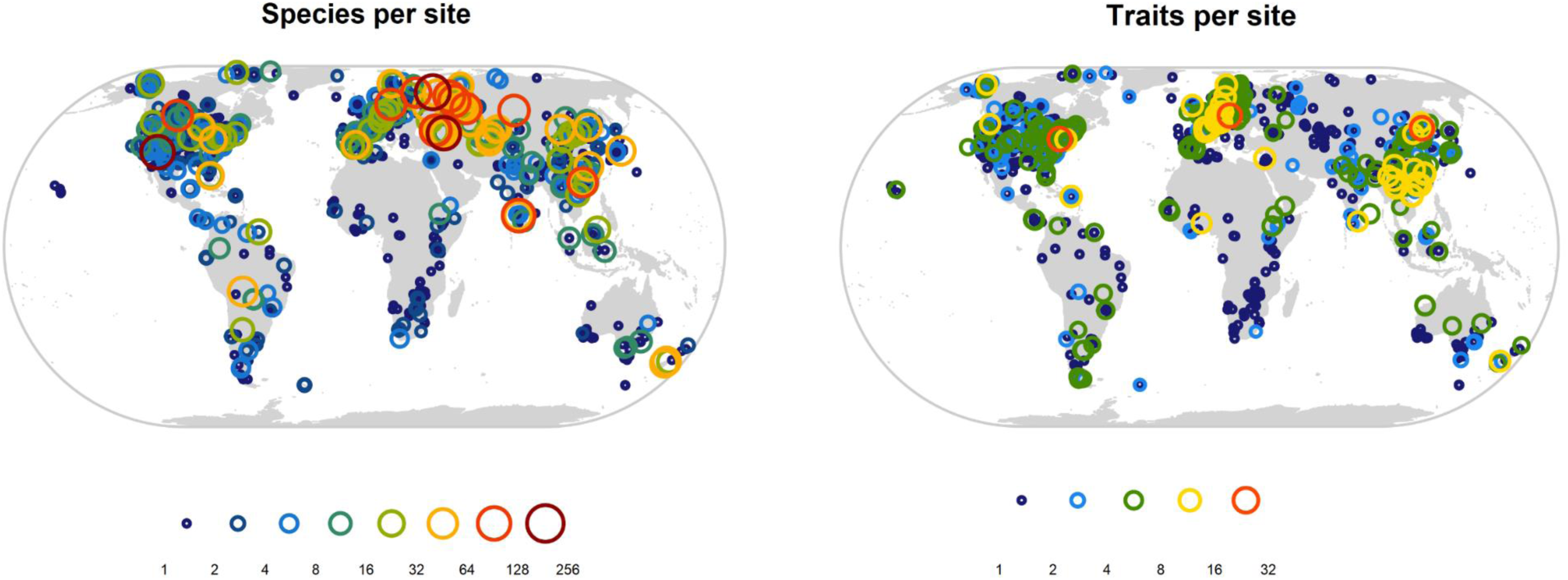
Maps depicting all geo-referenced data from field studies included in GRooT. Circles indicate range of species per site, e.g., 1-2 species, 2-4 species, successively, or trait per site, e.g., 1-2 traits, 2-4 traits successively.

Phylogenetically, data in GRooT cover all major clades of vascular plants, i.e., pteridophytes, gymnosperms, basal angiosperms, monocots, magnoliids, basal eudicots, superrosids and superasterids (Fig 2a), with data for 254 families. However, phylogenetic gaps are observed for traits related to key categories such as anatomy, architecture, dynamics, and physiology. When accounting for the number of vascular species included in GRooT (n = 6,214 species across 254 families), the average number of traits per species within family ranges between two to 14, with an overall average of four traits for species across the phylogeny. When accounting for the number of vascular species accepted globally (based on The Plant List; n = 316,110 species across n = 442 families), the average number of traits per species within family ranges from zero to eight traits, with an overall average of <1 traits for species (Fig 2b).

**Figure 2.**
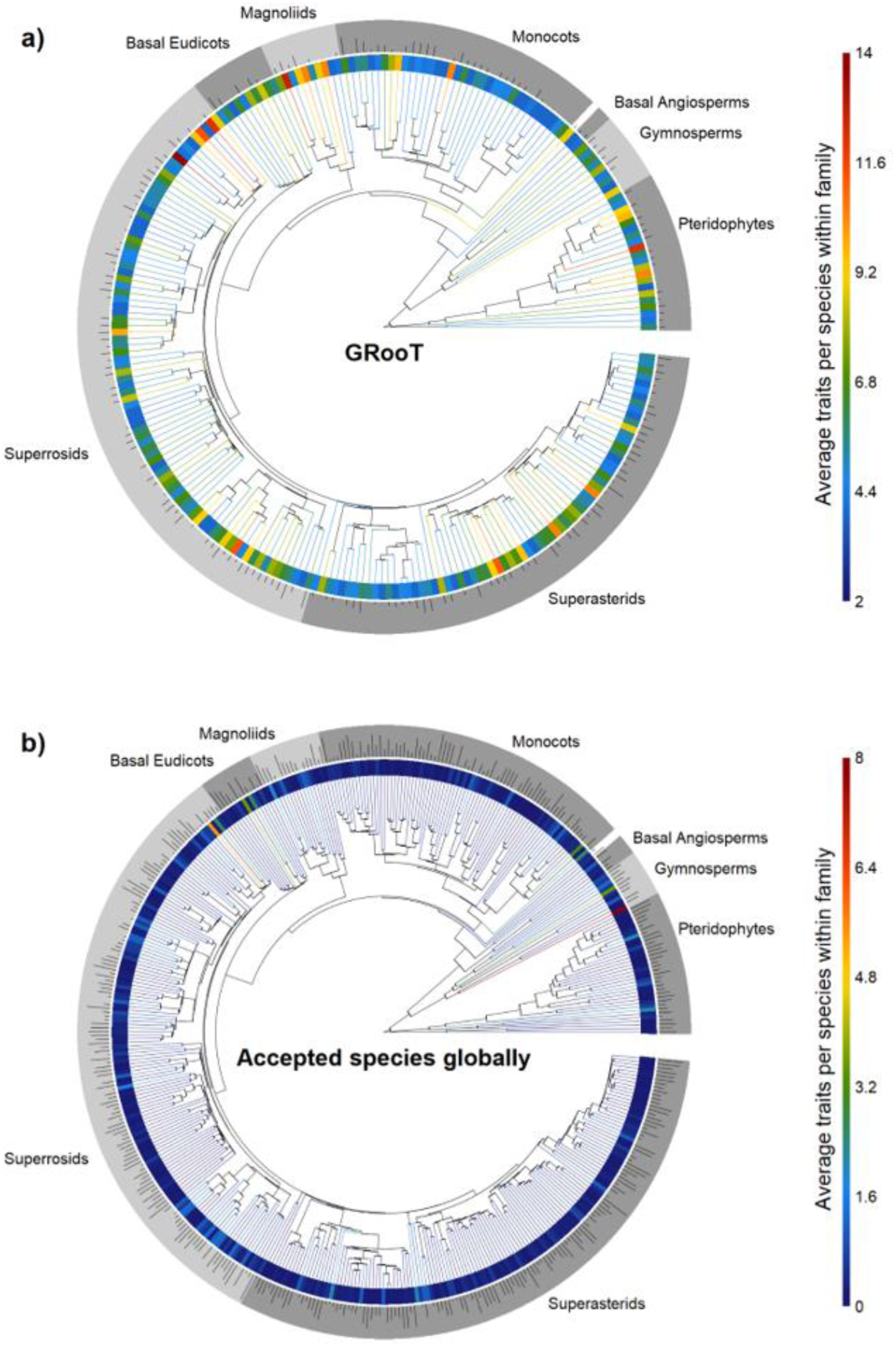
Phylogenetic coverage of root traits in GRoOT. Panel **(a)** shows the average distribution of root traits per species in GRooT across the phylogeny (n = 6,214 species across n = 254 families) and panel **(b)** shows GRooT phylogenetic coverage based on the accepted species by The Plant List (n = 316,110 species across n = 442 families). Tip and inner ring color depict mean number of traits per species in a family while dark blue color indicates families with lower number of traits per species. The outer ring represents major clades of vascular plants and the bars in this ring represent the family size (proportional to the logarithm base 10) either based on the number of species per family included in GRooT or the number of accepted species per family globally (Panel **a** and **b**, respectively).

## 4. Discussion

GRooT is a uniquely important step towards the inclusion of root traits in large-scale comparative studies and global models by integrating expert knowledge, data mobilization, standardization, curation and open accessibility. In terms of geo-referenced data from field studies, GRooT has highest coverage in North America, Europe, and Asia, especially for chemical and morphological traits, reflecting the capability for large-scale studies in these regions. In terms of phylogenetic coverage, data in GRooT include the major clades of vascular plants, with, on average, four traits included per species. Thereby, phylogenetic coverage in GRooT opens the possibility to use the data in large-scale phylogenetic studies like analyses of trait conservatism (Valverde-Barrantes *et al*., 2017; Averill *et al*., 2019) or assessments of trait relationships and trade-offs across the phylogeny.

GRooT also helps highlighting remaining barriers to integrate root trait data on global analyses. Particularly, data availability of certain relevant but hard-to-measure root traits related to physiology, mechanical properties and root dynamics generally remain scarce (Supplementary Information Table S2). Moreover, while GRooT contains global data with a wide geographical range, the species coverage in South America and Africa remains limited irrespective of trait type, reflecting overall biases in global ecological observations (Martin *et al*., 2012; Cornwell *et al*., 2019). Thus, targeted initiatives in these regions, such as that by Addo-Danso et al. (2019), are fundamental. While GRooT includes around 6,500 species, initiatives to increase the representativeness of species for families with highest species richness, such as Fabaceae, Fagaceae, Orchidaceae, and Poaceae, are also required.

GRooT can be used for – but is not restricted to – studying macroecological and functional biogeography (Violle *et al*., 2014), assessing global belowground trait-environmental relationships as known from aboveground approaches (Bruelheide *et al*., 2018), detecting fundamental ecological patterns such as the root economic space (Bergmann *et al*., 2020) or trade-offs and coordination among organs in the plant economic spectrum (Freschet *et al*., 2010). Furthermore, GRooT facilitates the integration of root traits into studies of related scientific disciplines, e.g. soil science and agronomy (Wood *et al*., 2015; Martin & Isaac, 2018). The completion of this standardized, curated and publicly-available database provide immediate benefit to the research community from ready-to-use data (Gallagher *et al*., 2019) and provide additional direction helping experts to identify gaps that need to be filled to increase completeness of global root trait data.

## Acknowledgments

This initiative was possible due to the support of the Synthesis Centre (sDiv) of the German Centre for Integrative Biodiversity Research (iDiv) Halle-Jena-Leipzig (DFG FZT 118) in the frame of the sRoot working group. We are grateful to all data contributors as well as to numerous assistants, technicians and students which made it possible to have these hard-earned valuable data in the first place. We thank Owen Atkin, F Stuart Chapin III, Brian Enquist, Thomas Hickler, Belinda Medlyn, Quentin Read, Michael White, and Christian Wirth for contributing data, Jitka Klimesová for contributing data and providing comments in an earlier version of the manuscript, and A. Shafer Powell for assistance with FRED user guidance and data. We acknowledge the ECOCRAFT data contributors: Ana Rey, Craig Barton, Gail Jackson, Bernard Saugier, Franz Badeck, Eric Dufrêne, Rodolphe Liozon, Dieter Overdieck, Manfred Forstreuter, Joern Strassemeyer, Andrew Friend, Maureen Murray, Giuseppe Scarascia-Mugnozza, Giorgio Matteuci, Paolo de Angelis, Mark Broadmeadow, Tim Randle, Eric Laitat, Bruno Portier, Bruno Chermanne, Reinhart Ceulemans, David de Pury, Ewa Jach, Ivan Janssens, Sune Linder, Johan Bergh, Sabine Philippot, Peter Roberntz, Jan Stockfors, Seppo Kellomaki, Kaisa Laitinen, Kaiyun Wang, Michal Marek, and Breta Regner. Colleen M. Iversen and the Fine-Root Ecology Database (FRED) are supported by the Biological and Environmental Research program in the Department of Energy’s Office of Science and Joana Bergmann was supported by DFG grants RU-1815/20-1 and RI 1815/22-1. Nathaly Guerrero-Ramírez thanks the Dorothea Schlözer Postdoctoral Programme of the Georg-August-Universität Goettingen for their support, Nadejda A. Soudzilovskaia thanks vidi grant 016.161.318 issued by the Netherlands Organization for Scientific Research and Vladimir G. Onipchenko thanks RSF for financial support (#16-14-10208). The study has been supported by the TRY initiative on plant traits (http://www.try-db.org). The TRY initiative and database is hosted, developed and maintained at the Max Planck Institute for Biogeochemistry, Jena, Germany. TRY is currently supported by DIVERSITAS/Future Earth and the German Centre for Integrative Biodiversity Research (iDiv) Halle-Jena-Leipzig.

## Biosketch

**The Global Root Trait Research Team** is a group of root ecologist interested on contributing towards the inclusion of root traits in large-scale comparative studies and global models by offering standardized and publicly curated data of key root traits. The team built GRooT during two synthesis workshops on root traits (sRoot working group) and with the help of external researchers. We have also developed interconnections with other databases, creating innovative linkages and facilitating the use of complementary information among databases. Our goal is to provide accessible information to overcome the conceptual and methodological roadblocks limiting the use of root traits by a wide community of ecologist and biogeographers assessing topics such as global belowground trait-environmental relationships, detecting ecological patterns such as the root economic space or trade-offs and coordination among organs in the plant economic spectrum.

## Data availability

GRooT is publicly available, the database csv files and the script in R to query the database are stored in a GitHub repository (https://github.com/GRooT-Database/GRooT-Data).

GitHub page: (https://groot-database.github.io/GRooT/).

## Supporting Information

**Table S1.** Definitions and units of root traits included in GRooT. Definitions based on the handbook of root traits (Freschet *et al*., submitted), FRED guidelines (Iversen *et al*., 2018) and CLO-PLA (Klimešová & Bello, 2009; Klimešová *et al*., 2017, 2019). Traits categorization based on the handbook of root traits and McCormack et al (2017).

| Traits | Units | Definitions |
| --- | --- | --- |
| **Anatomy** |  |  |
| Root cortex thickness | μm | Thickness of the ring of cortical cells beginning outside the stele and extending to the root epidermis |
| Root stele diameter | μm | Diameter of root stele |
| Root stele fraction | % | Root cross-sectional area that is occupied by the stele |
| Root xylem vessel number | number mm<sup>-2</sup> | Root xylem vessel number per root stele area |
| Root vessel diameter | μm | Root vessel diameter |
| **Architecture** |  |  |
| Root branching ratio | number number<sup>-1</sup> | Number of roots on a given order divided by the number of roots in the higher order |
| Root branching density | number cm<sup>-1</sup> | Number of laterals on a given length unit of total root length<sup>t</sup> |
| **Belowground allocation** |  |  |
| Root mass fraction | g g<sup>-1</sup> | Ratio of root dry mass per total plant standing dry biomass |
| **Chemistry** |  |  |
| Root structural carbohydrate concentration | mg g<sup>-1</sup> | Root cellulose and hemicellulose content per root mass |
| Root lignin concentration | mg g<sup>-1</sup> | Mass of lignin per root dry mass |
| Root carbon concentration | mg g<sup>-1</sup> | Mass of C per root dry mass |
| Root nitrogen concentration | mg g<sup>-1</sup> | Mass of N per root dry mass |
| Root phosphorus concentration | mg g<sup>-1</sup> | Mass of P per root dry mass |
| Root carbon to nitrogen ratio | g g<sup>-1</sup> | Ratio of carbon to nitrogen in root by mass |
| Root nitrogen to phosphorus ratio | g g<sup>-1</sup> | Ratio of nitrogen to phosphorus in root by mass |
| Root calcium concentration | mg g<sup>-1</sup> | Mass of Ca per root dry mass |
| Root potassium concentration | mg g<sup>-1</sup> | Mass of K per root dry mass |
| Root magnesium concentration | mg g<sup>-1</sup> | Mass of Mg per root dry mass |
| Root manganese concentration | mg g<sup>-1</sup> | Mass of Mn per root dry mass |
| ***Dynamics & decomposition*** |  |  |
| Root production | g m<sup>-2</sup> yr<sup>-1</sup> | Dry mass of root produced per soil volume and per year |
| Root lifespan | days | Time between birth and death of a root |
| Root turnover rate | year<sup>-1</sup> | Root dry mass production per dry mass of a given pool of roots over a period of time |
| Root litter mass loss rate | year<sup>-1</sup> | Dry mass lost by roots per initial dry mass of the roots per unit time |
| ***Horizontal plant mobility*** |  |  |
| Lateral spread | cm yr<sup>-1</sup> | Distance a clonal plant grows laterally in one year |
| ***Microbial associations*** |  |  |
| Root mycorrhizal colonization intensity | % | Percentage length of roots colonized by fungi |
| ***Morphology*** |  |  |
| Mean root diameter | mm | Mean diameter of the root sample |
| Root dry matter content | g g<sup>-1</sup> | Dry mass of root per unit fresh root mass |
| Root tissue density | g cm<sup>-3</sup> | Dry mass of root per unit volume of fresh root |
| Specific root area | cm<sup>2</sup> g<sup>-1</sup> | Surface area of a root per unit dry mass |
| Specific root length | m g<sup>-1</sup> | Length of a root per unit dry mass |
| ***Physiology & respiration*** |  |  |
| Net uptake rate of nitrogen | μmol g<sup>-1</sup> d<sup>-1</sup> | Amount of labelled nitrogen accumulated within a plant individual per unit dry mass and time<sup>†</sup> |
| Specific root respiration | nmol g<sup>-1</sup> s<sup>-1</sup> | Amount of CO<sub>2</sub> released or O<sub>2</sub> absorbed by root per unit root dry mass per unit of time |
| ***System & distribution*** |  |  |
| Fine root mass to leaf mass ratio | g g<sup>-1</sup> | Ratio of fine-root mass to leaf mass |
| Coarse to fine root mass ratio | g g<sup>-1</sup> | Ratio of coarse root biomass to fine root biomass |
| Root mass density | g cm<sup>-3</sup> | Mass of roots per unit soil volume |
| Root length density | cm cm<sup>-3</sup> | Length of roots per unit soil volume |
| Maximum rooting depth | m | Maximum soil depth at which roots occur |

<sup>†</sup> As the total root length in the category can vary across studies, this information needs to be interpreted with caution.
<sup>‡</sup> Combined uptake of <sup>15</sup>N as NO<sub>3</sub><sup>-</sup>, NH<sub>4</sub><sup>+</sup> or glycine tracers in 24h per shoot dry mass, estimated as $F = [T(AS-AB)]/AF$ with T is the plant N concentration, AS is the atom percent excess <sup>15</sup>N in the sample, AB is the atom percent excess <sup>15</sup>N in natural sample, AF is the atom percent excess <sup>15</sup>N in the tracer

**Table S2.** Relevant root traits not included in GRooT due to low coverage (data for <50 species available). Definitions based on the handbook of root traits (Freschet *et al*., submitted) and FRED guidelines (Iversen et al. 2018). Traits categorization based on the handbook of root traits and McCormack et al. (2017).

| Traits | Units | Definitions |
| --- | --- | --- |
| **Architecture** |  |  |
| Root branching intensity | number cm<sup>-1</sup> | number of laterals on a given length unit of parent root |
| **Chemistry** |  |  |
| Root total non-structural carbohydrate concentration | mg g<sup>-1</sup> | the mass of carbohydrate molecules that do not participate in root structure per root dry mass |
| Root phenolic concentration | mg g<sup>-1</sup> | the mass of phenolic compounds per root dry mass |
| Root sulphur (S) concentration | mg g<sup>-1</sup> | the mass of S per root dry mass |
| Root aluminum concentration | mg g<sup>-1</sup> | the mass of Al per root dry mass |
| **Dynamics** |  |  |
| Time root growth initiation | Date | the calendar date at which roots production stops or becomes minimal after a period of high growth |
| Time root growth cessation | Date | the calendar date at which root production starts increasing after a period of low or null growth |
| **Mechanical** |  |  |
| Root tensile strength | Pascal | The force required to cause failure of the root per cross-sectional area |
| **Morphology** |  |  |
| Root hair density | number mm<sup>-2</sup> | the number of root hairs per unit root length |
| Root hair length | µm | the length of fully-grown root hairs |
| **Physiology** |  |  |
| Theoretical root specific hydraulic conductance | m<sup>3</sup> s<sup>-1</sup> MPa<sup>-1</sup> cm<sup>-2</sup> | the calculated amount of water that can move through tissue per the cross sectional area of the tissue |
| Maximum net ion uptake rate, I<sub>max</sub> | µmol g<sup>-1</sup>s<sup>-1</sup> | the amount of ion accumulated per unit root biomass and time under conditions of non-limiting nutrient concentration |
| Michaelis-Menten constant, km | mmol l<sup>-1</sup> or mM | the nutrient concentration where 50% of the maximum net ion uptake rate is observed |
| **Root system** |  |  |
| Vertical root mass distribution index | Unitless | Extinction coefficient (β) of an asymptotic equation (Y = 1 - β<sup>d</sup>) fitting the cumulative proportional root biomass over depth. Values of β approaching 1 imply a greater proportion of roots with depth |
| Lateral root mass distribution index | Unitless | Extinction coefficient (β) of an asymptotic equation (Y = 1 - β<sup>d</sup>) fitting the cumulative proportional root biomass over depth. Values of β approaching 1 correspond to a greater proportion of roots away from the plant base |
| Lateral rooting extent | m | Maximum distance between superficial roots and the base of the plant |

**Table S3.**
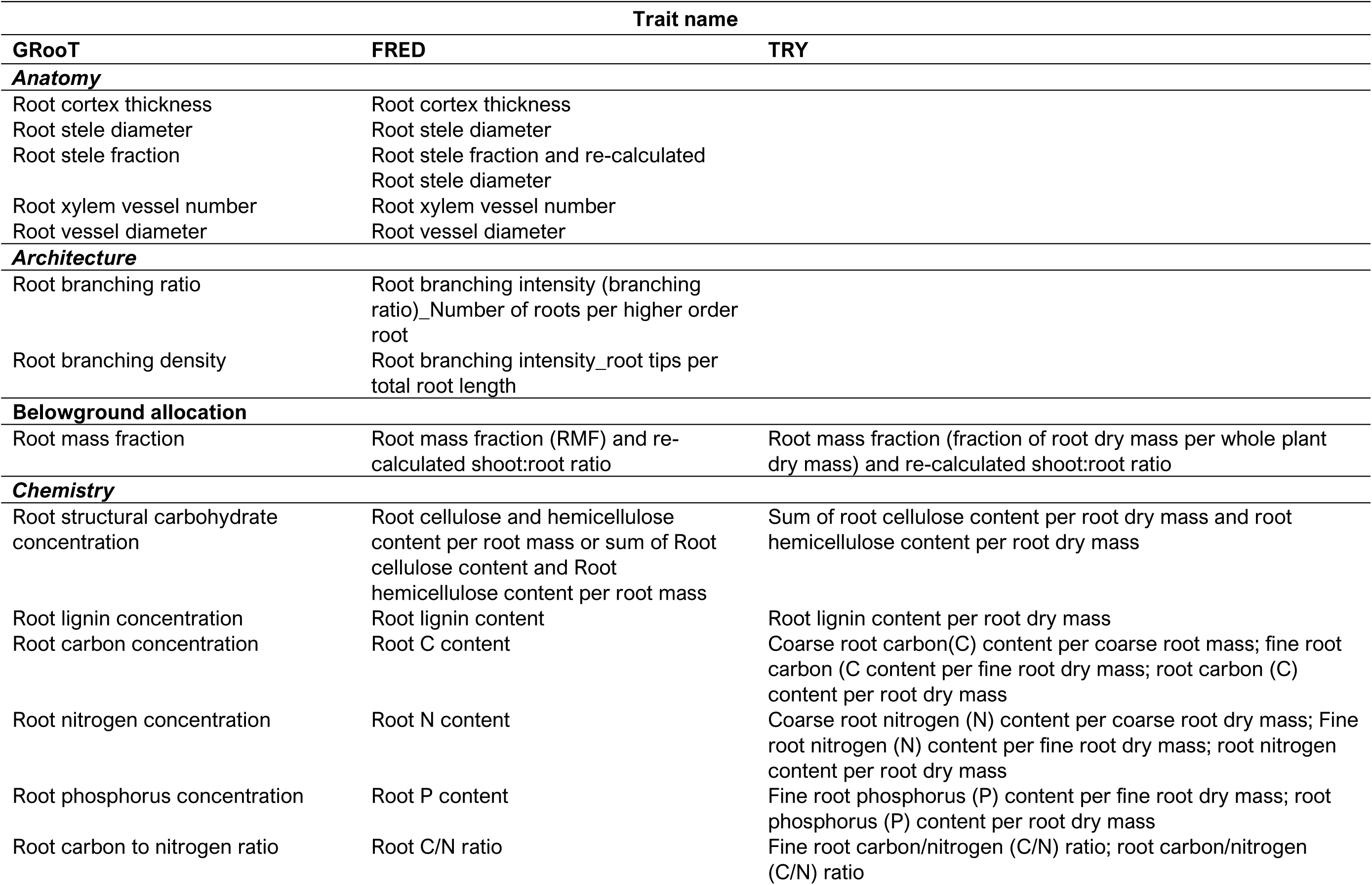
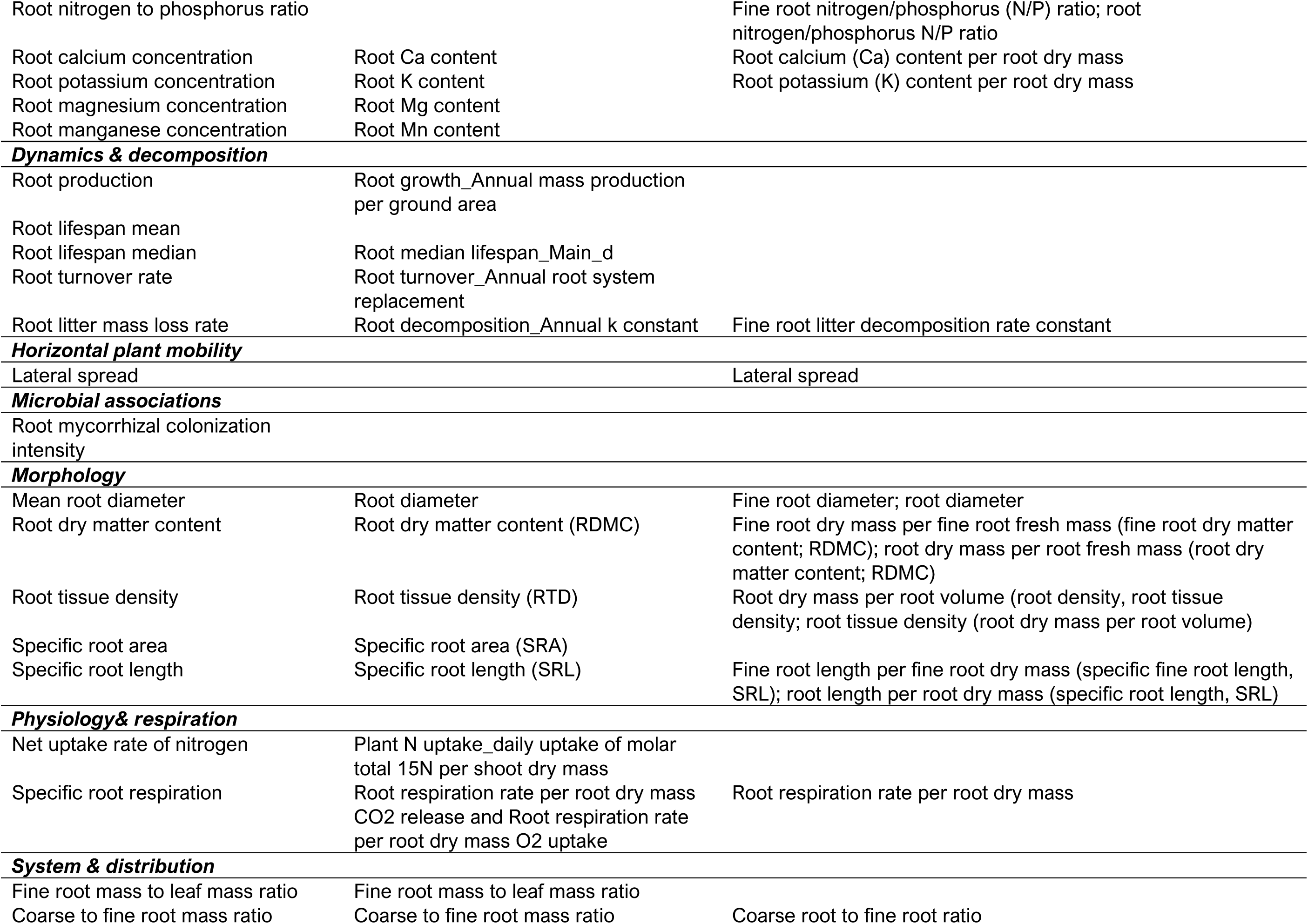
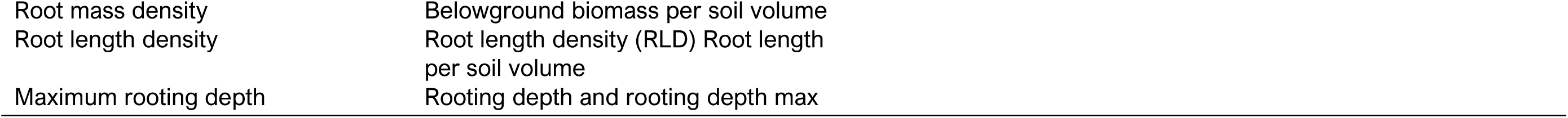
Trait names across major trait databases with root traits. FRED is also included in the TRY Database. Aggregable trait variables into a single unique trait in GRooT are showed.

| Trait name |  |  |
| --- | --- | --- |
| GRooT | FRED | TRY |
| **Anatomy** |  |  |
| Root cortex thickness | Root cortex thickness |  |
| Root stele diameter | Root stele diameter |  |
| Root stele fraction | Root stele fraction and re-calculated<br>Root stele diameter |  |
| Root xylem vessel number | Root xylem vessel number |  |
| Root vessel diameter | Root vessel diameter |  |
| **Architecture** |  |  |
| Root branching ratio | Root branching intensity (branching<br>ratio)_Number of roots per higher order<br>root |  |
| Root branching density | Root branching intensity_root tips per<br>total root length |  |
| **Belowground allocation** |  |  |
| Root mass fraction | Root mass fraction (RMF) and re-calculated shoot:root ratio | Root mass fraction (fraction of root dry mass per whole plant<br>dry mass) and re-calculated shoot:root ratio |
| **Chemistry** |  |  |
| Root structural carbohydrate<br>concentration | Root cellulose and hemicellulose<br>content per root mass or sum of Root<br>cellulose content and Root<br>hemicellulose content per root mass | Sum of root cellulose content per root dry mass and root<br>hemicellulose content per root dry mass |
| Root lignin concentration | Root lignin content | Root lignin content per root dry mass |
| Root carbon concentration | Root C content | Coarse root carbon(C) content per coarse root mass; fine root<br>carbon (C content per fine root dry mass; root carbon (C)<br>content per root dry mass |
| Root nitrogen concentration | Root N content | Coarse root nitrogen (N) content per coarse root dry mass; Fine<br>root nitrogen (N) content per fine root dry mass; root nitrogen<br>content per root dry mass |
| Root phosphorus concentration | Root P content | Fine root phosphorus (P) content per fine root dry mass; root<br>phosphorus (P) content per root dry mass |
| Root carbon to nitrogen ratio | Root C/N ratio | Fine root carbon/nitrogen (C/N) ratio; root carbon/nitrogen<br>(C/N) ratio |
| Root nitrogen to phosphorus ratio |  | Fine root nitrogen/phosphorus (N/P) ratio; root nitrogen/phosphorus N/P ratio |
| Root calcium concentration | Root Ca content | Root calcium (Ca) content per root dry mass |
| Root potassium concentration | Root K content | Root potassium (K) content per root dry mass |
| Root magnesium concentration | Root Mg content |  |
| Root manganese concentration | Root Mn content |  |
| **Dynamics & decomposition** |  |  |
| Root production | Root growth_Annual mass production per ground area |  |
| Root lifespan mean |  |  |
| Root lifespan median | Root median lifespan_Main_d |  |
| Root turnover rate | Root turnover_Annual root system replacement |  |
| Root litter mass loss rate | Root decomposition_Annual k constant | Fine root litter decomposition rate constant |
| **Horizontal plant mobility** |  |  |
| Lateral spread |  | Lateral spread |
| **Microbial associations** |  |  |
| Root mycorrhizal colonization intensity |  |  |
| **Morphology** |  |  |
| Mean root diameter | Root diameter | Fine root diameter; root diameter |
| Root dry matter content | Root dry matter content (RDMC) | Fine root dry mass per fine root fresh mass (fine root dry matter content; RDMC); root dry mass per root fresh mass (root dry matter content; RDMC) |
| Root tissue density | Root tissue density (RTD) | Root dry mass per root volume (root density, root tissue density; root tissue density (root dry mass per root volume)) |
| Specific root area | Specific root area (SRA) |  |
| Specific root length | Specific root length (SRL) | Fine root length per fine root dry mass (specific fine root length, SRL); root length per root dry mass (specific root length, SRL) |
| **Physiology& respiration** |  |  |
| Net uptake rate of nitrogen | Plant N uptake_daily uptake of molar total 15N per shoot dry mass |  |
| Specific root respiration | Root respiration rate per root dry mass CO2 release and Root respiration rate per root dry mass O2 uptake | Root respiration rate per root dry mass |
| **System & distribution** |  |  |
| Fine root mass to leaf mass ratio | Fine root mass to leaf mass ratio |  |
| Coarse to fine root mass ratio | Coarse to fine root mass ratio | Coarse root to fine root ratio |

|  |  |
| --- | --- |
| Root mass density | Belowground biomass per soil volume |
| Root length density | Root length density (RLD) Root length<br>per soil volume |
| Maximum rooting depth | Rooting depth and rooting depth max |

**Table S4.** Available information on root entities available in GRooT. Definitions from the handbook of root traits (Freschet *et al*., submitted) and FRED guidelines (Iversen *et al*., 2018).

| Column name | Definition | Categories and definitions |
| --- | --- | --- |
| belowgroundEntities<sup>†</sup> | Selected main entities | *coarse roots (CR)*: root with large diameter, often roots > 2 mm in diameter; roots are generally lignified, with clear secondary development<br>*fine roots (FR)*: root with small diameter, often roots ≤ 2 mm in diameter; roots are generally not lignified<br>*total root system (TR)*: roots considered without specifying if coarse roots are included<br>*total belowground (TB)*: belowground parts included but not specified<br>*belowground steam (BS)<sup>‡</sup>*: belowground steams<br>*rhizomes (R)<sup>‡</sup>*: usually shoot axis growing horizontally at or below the substrate and produces shoots above and adventitious roots below<br>*unspecified*: information about entities not provided |
| belowgroundEntitiesOrderClassification | Root order-based classifications based on root branch hierarchy | *centrifugal*: also knows as developmental classification, it is a root- (i.e. growth axis-) based approach in which the lowest order root is the basal or shoot-borne root (i.e. order '0') and the highest order roots are the most distal (coarsest to finest)<br>*centripetal*: also knows as morphometric or stream-order ('Strahler') classification, it is a segment order-based approach in which distal root segments are first-order and parent root segments are higher order (finest to coarsest) |
| belowgroundEntitiesOrder | Root branching order, either coarsest to finest (Centrifugal) or finest to coarsest (Centripetal). | numeric |
| belowgroundEntitiesOrderMin | Minimum root order included | numeric |
| belowgroundEntitiesOrderMax | Maximum root order included | numeric |
| belowgroundEntitiesFuctional | Root functional classification based on functionally similar categories of absorptive and transport roots. The absorptive/transport function is associated mostly with the presence/absence of phellem | *absorptive<sup>\*</sup>*: root with dominantly absorptive function<br>*transport<sup>\*</sup>*: root with dominantly transport function |
| BelowgroundEntitiesDiameterMin** | Minimum root diameter included | numeric |
| BelowgroundEntitiesDiameterMax** | Maximum root diameter included | numeric |

†Multiple categories together are also included
‡Belowground stem (BS) and rhizomes (R) were only included when measured together with roots.
\*Both categories together are also included
\*\*Information of root diameter in the traitName column can be used too for determining entities based on diameter cut-offs

**Table S5.**
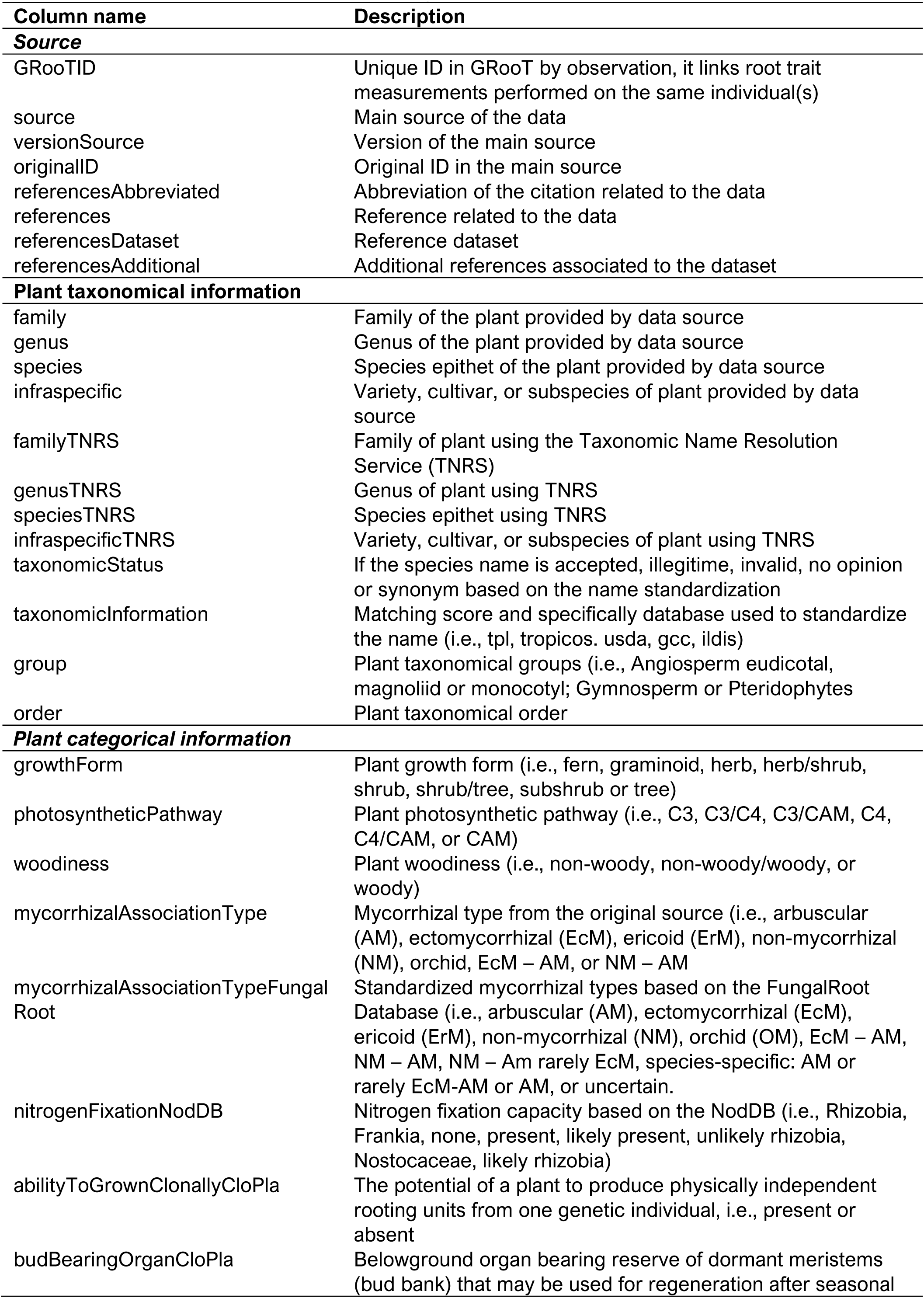
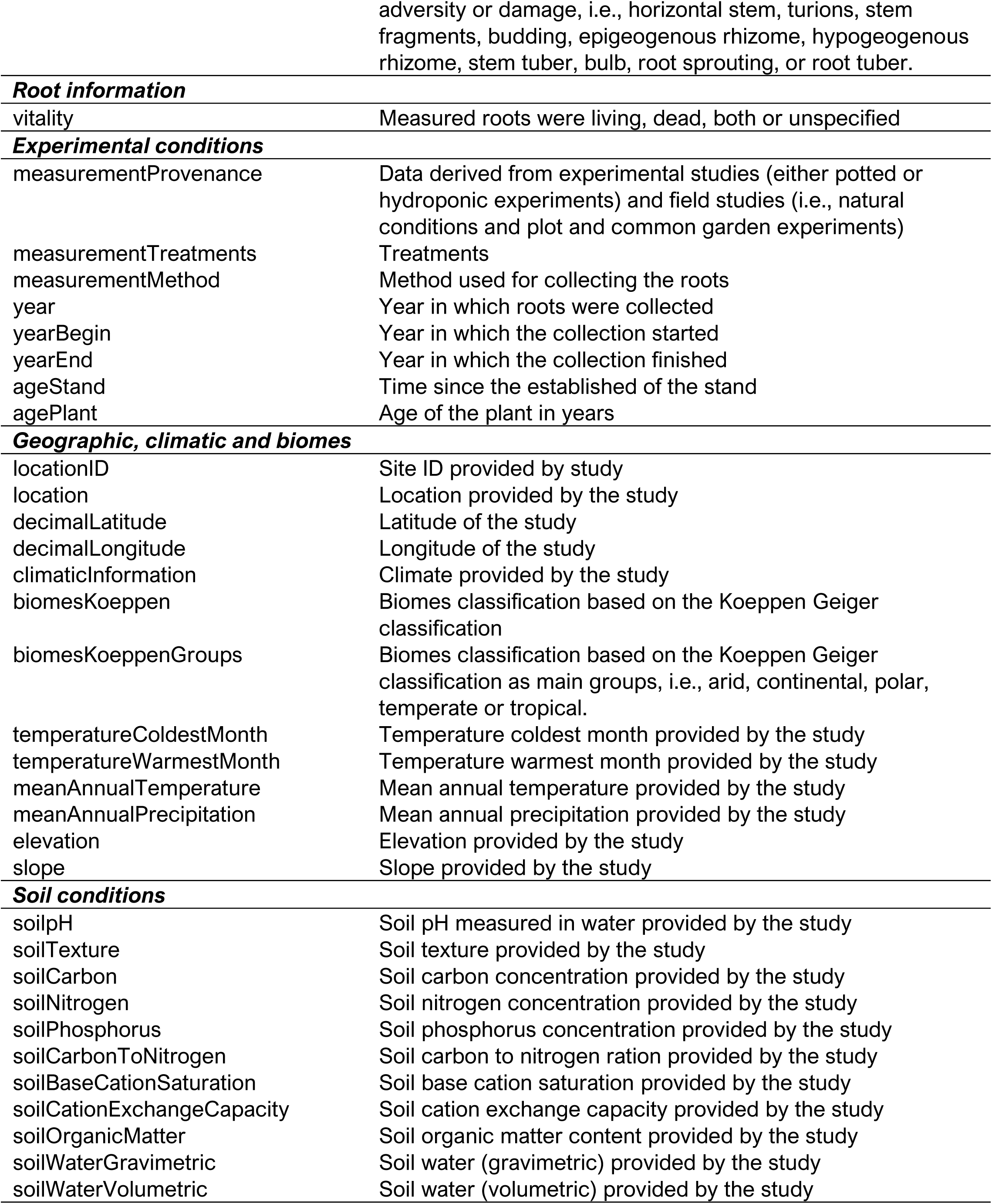
Selected information and meta-data provided in GRooT.

### Standardization across traits

Root mass fraction (RMF) was calculated from data of root-to-shoot biomass ratio (R:S) by:

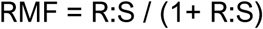

Root stele fraction was calculated using information on root stele diameter and root diameter.

Data from papers reporting stele fraction, stele diameter and root diameter were used to compare between stele fraction measured directly and calculated values (Fig S1)

**Figure S1.**
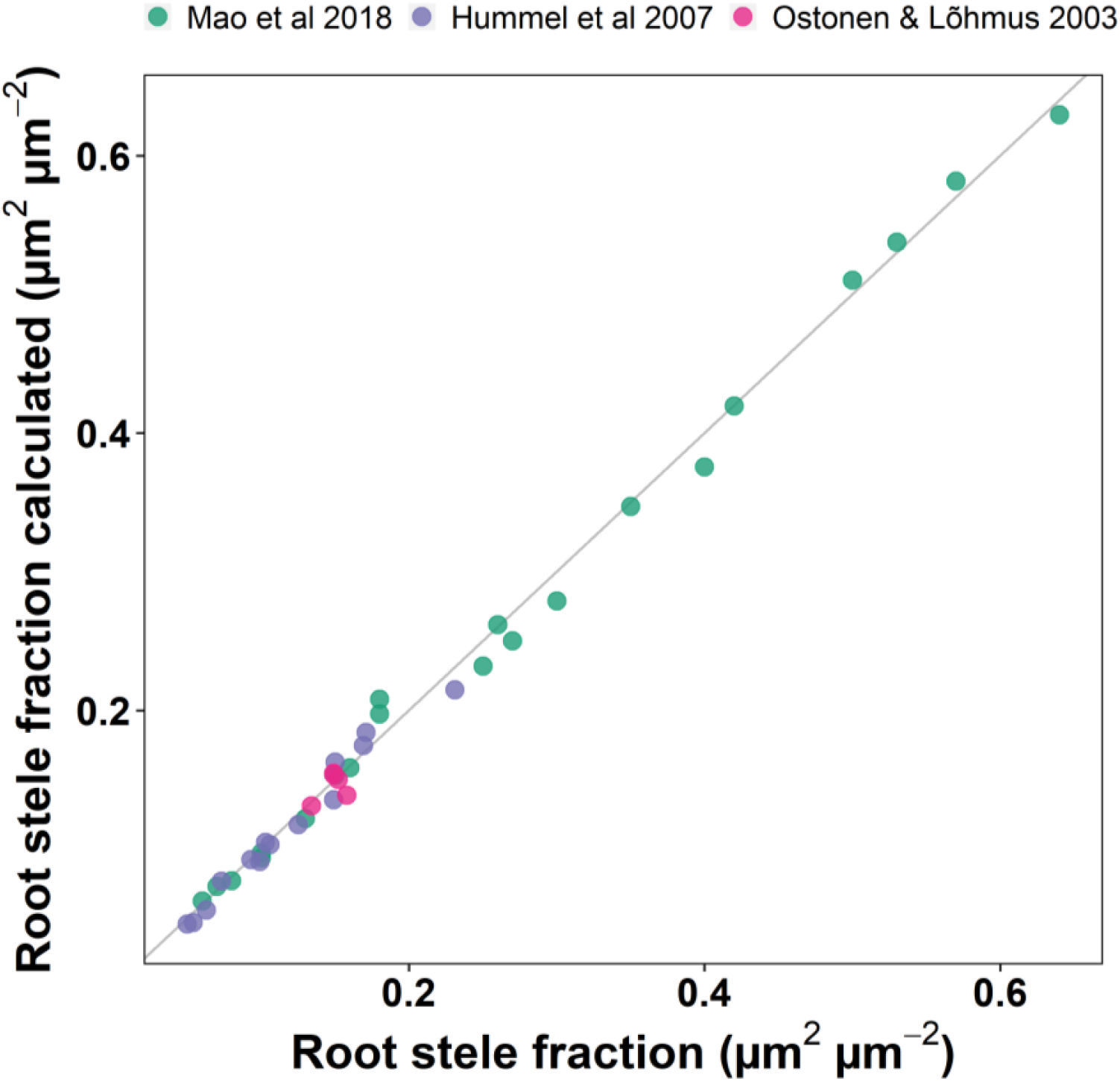
Comparison between root stele fraction reported (based on cross sectional area directly measured; x axis) and the calculated root stele fraction (y axis), determined by calculating cross sectional area of stele and root based on stele and root diameter, respectively. Points show studies in the database, that have data for both, area and diameter. The gray line has an intercept of 0 and a slope of 1. Results from a standardized major axis estimation show a positive relationship between the calculated and measured root stele fractions (R^2^: 0.99, p-value 2.22e^-16^).

**Figure S2.**
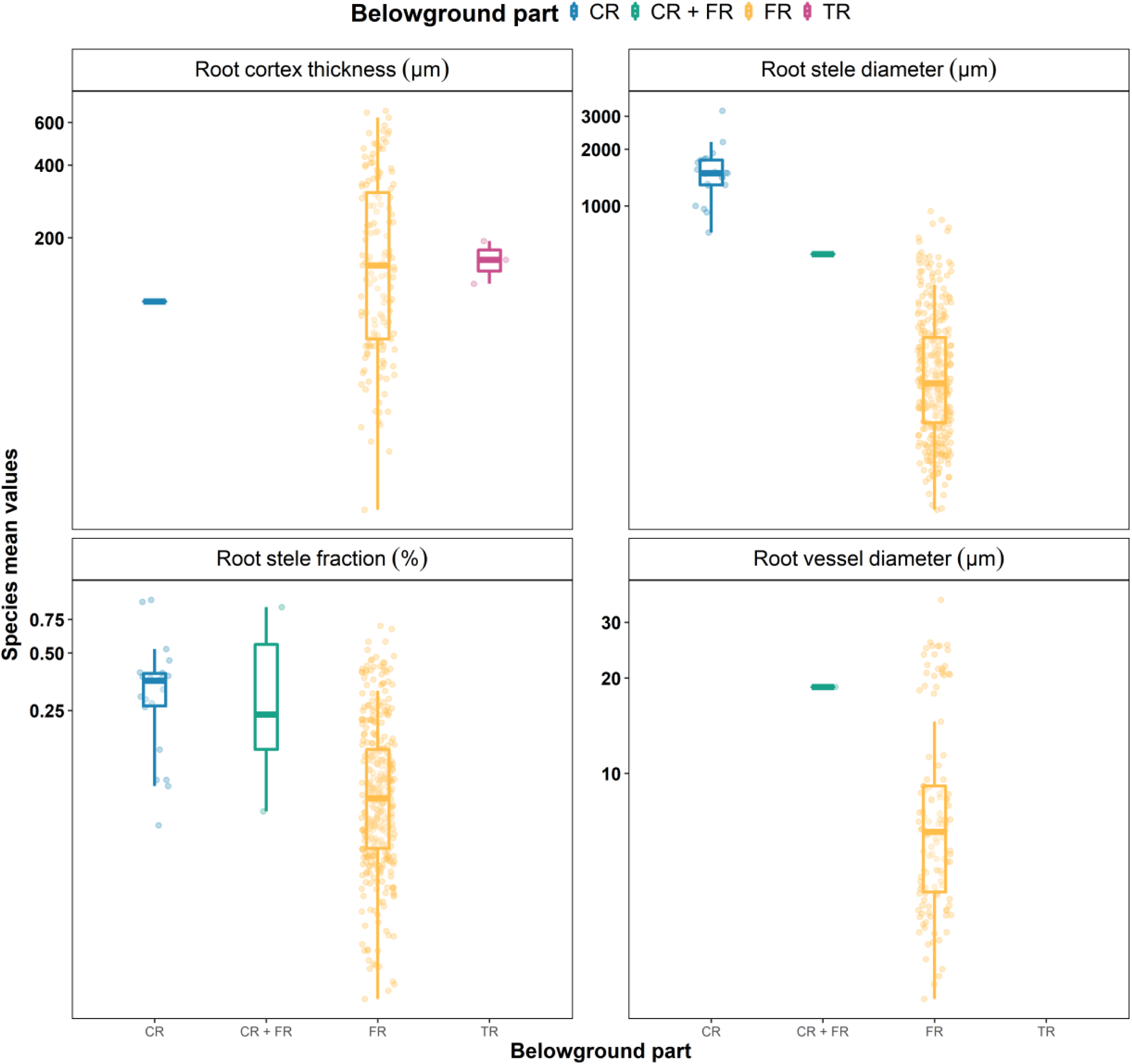
Density plots for anatomical traits. Points represent species mean values. Axis “y” is logarithmic base 2. Data from coarse (CR), fine (FR) and total roots (TR).

**Figure S3.**
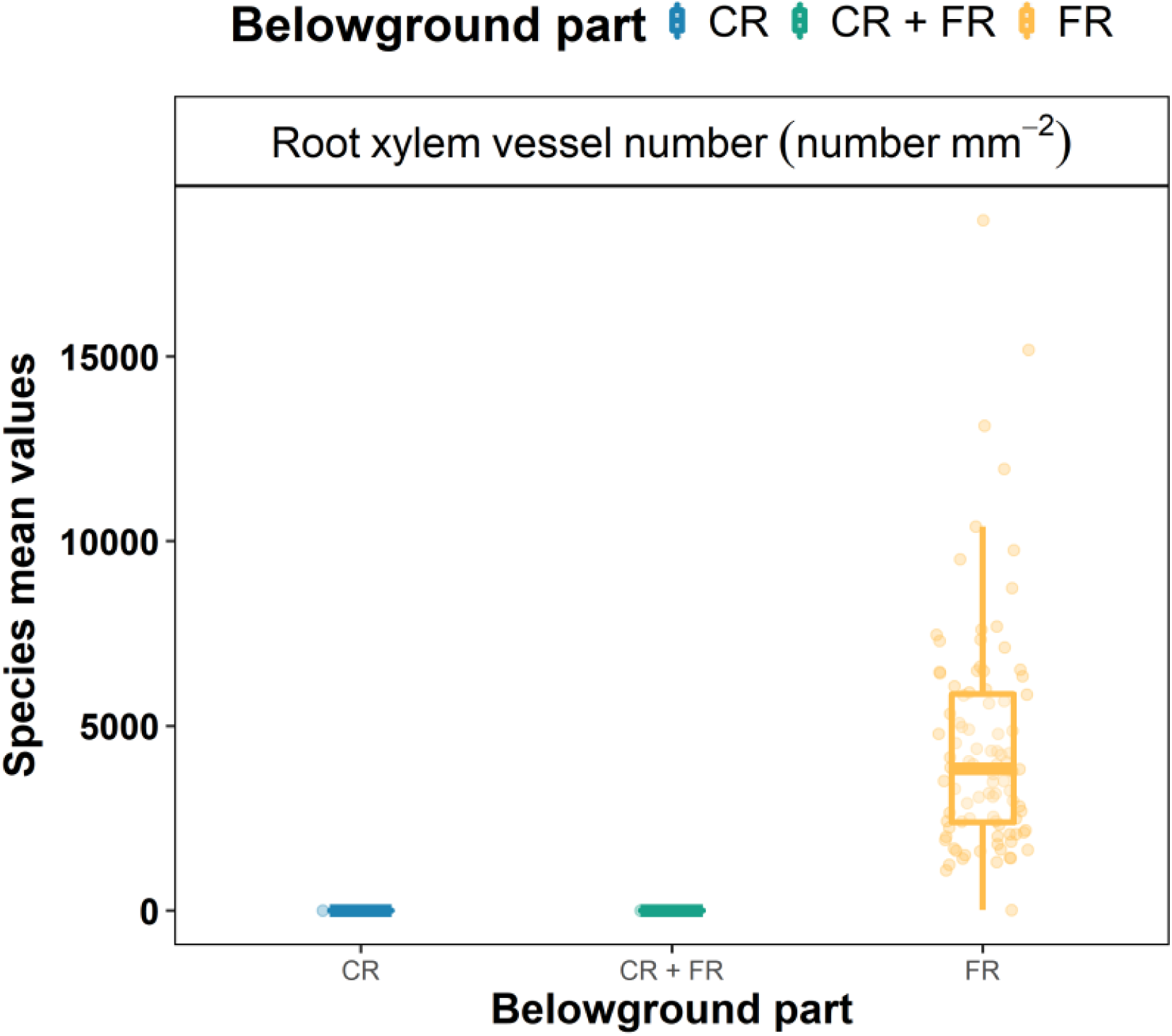
Density plots for root xylem vessel number. Points represent species mean values. Data from coarse (CR) and fine roots (FR).

**Figure S4.**
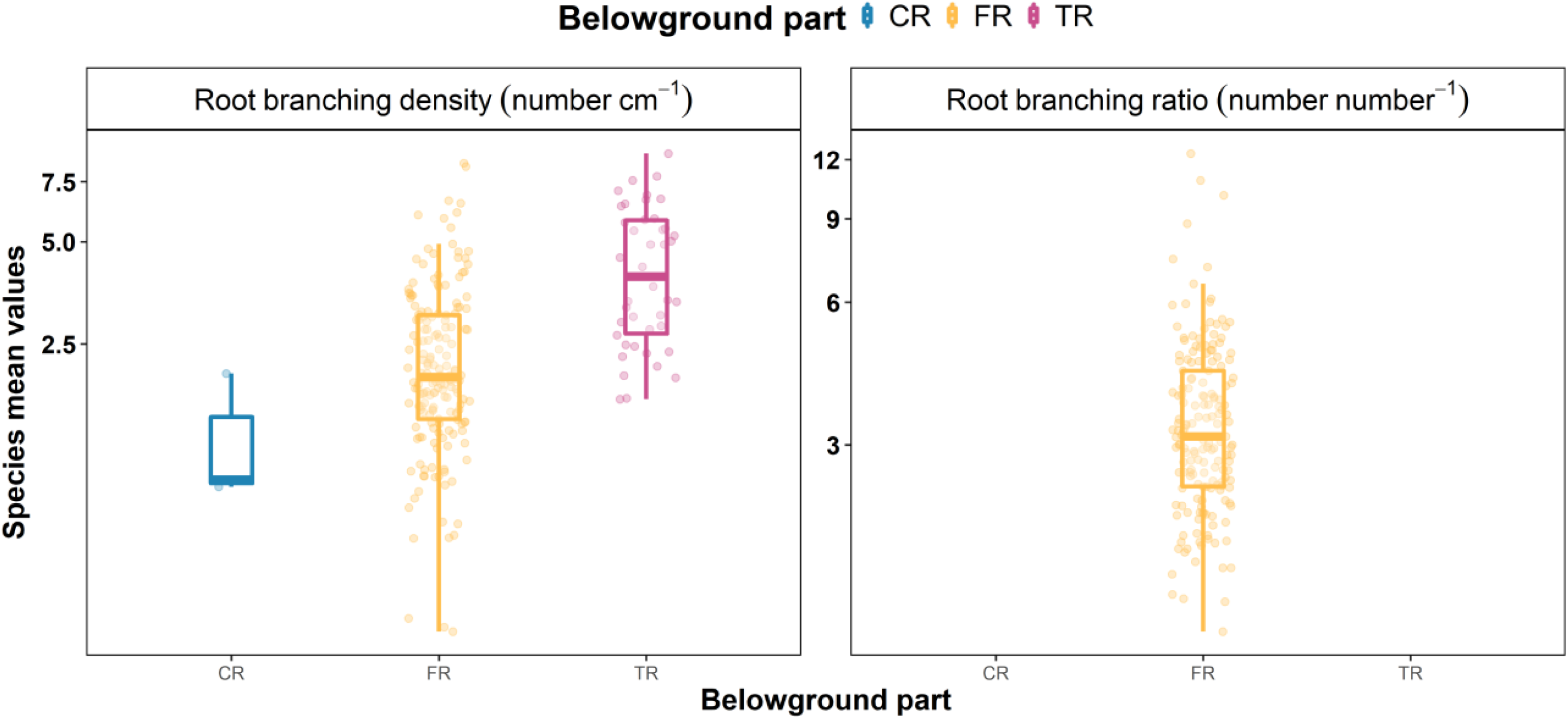
Density plots for architectural traits. Points represent species mean values. Axis “y” is logarithmic base 2. Data from coarse (CR), fine (FR) and total roots (TR).

**Figure S5.**
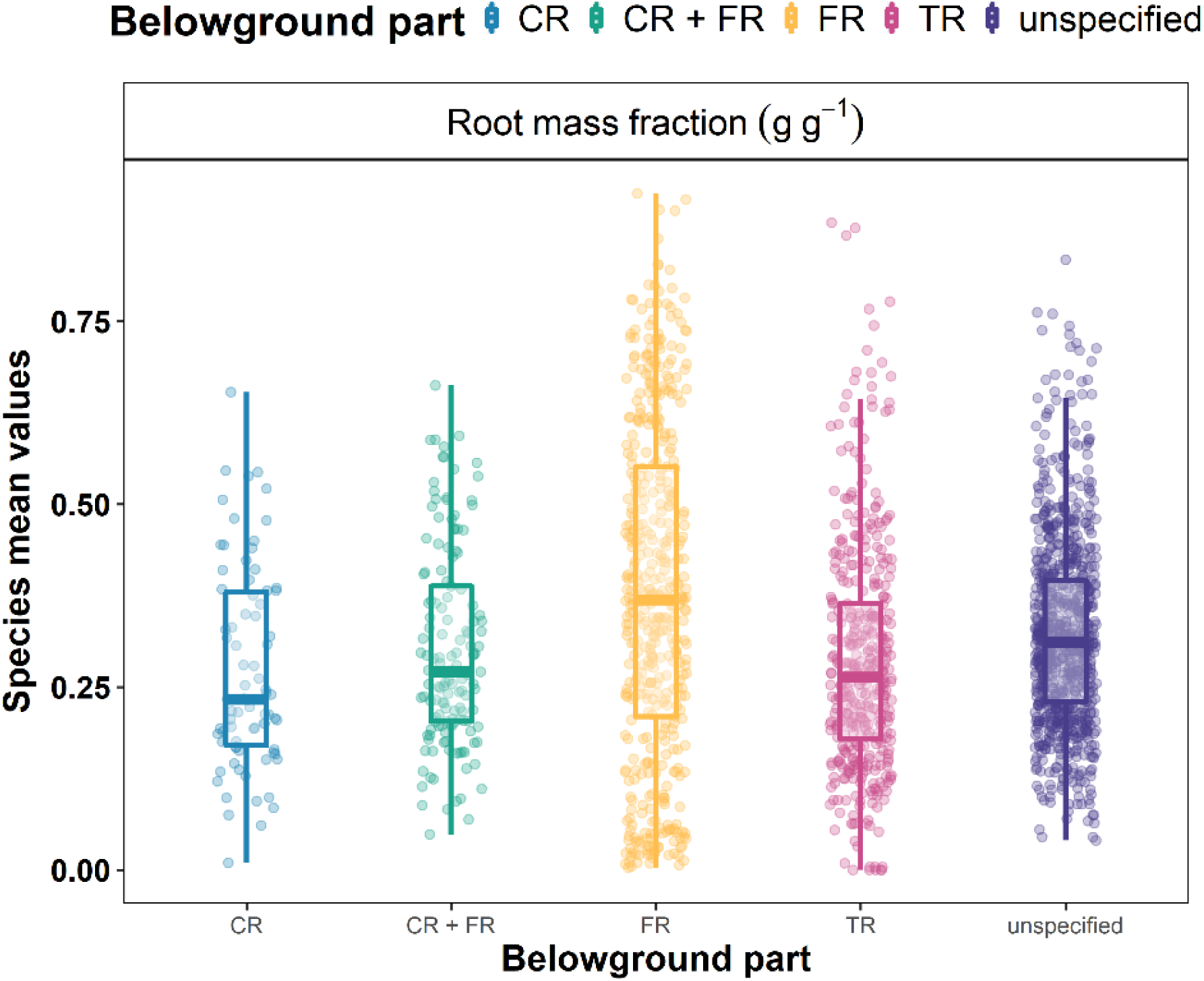
Density plots for root mass fraction. Points represent species mean values. Data from coarse (CR), fine (FR) and total roots (TR) or unspecified.

**Figure S6.**
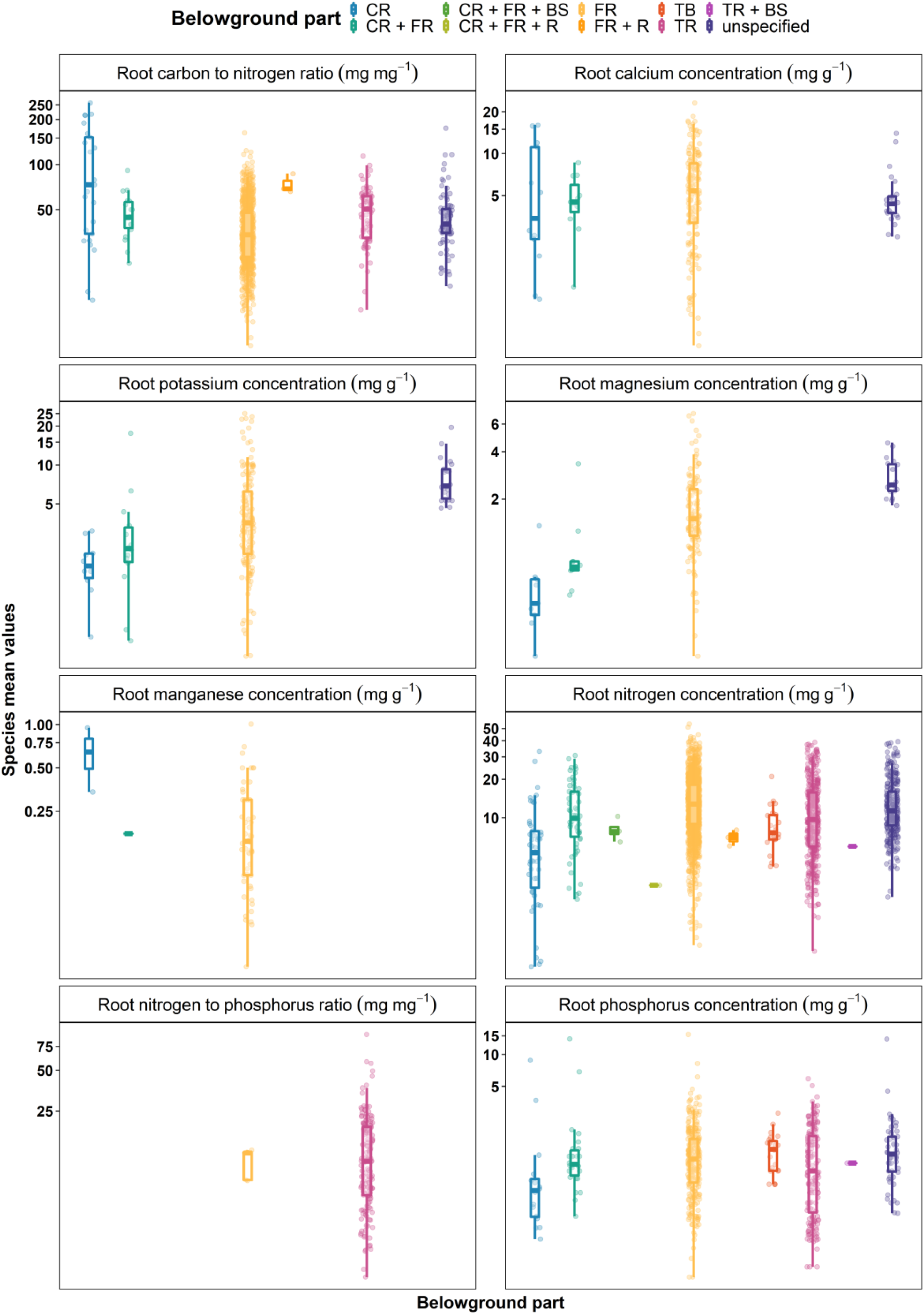
Density plots for chemical traits. Points represent species mean values. Axis “y” is logarithmic base 2. Data from coarse (CR), fine (FR) and total roots (TR), belowground steam (BS), rhizomes (R), total belowground (TB) or unspecified.

**Figure S7.**
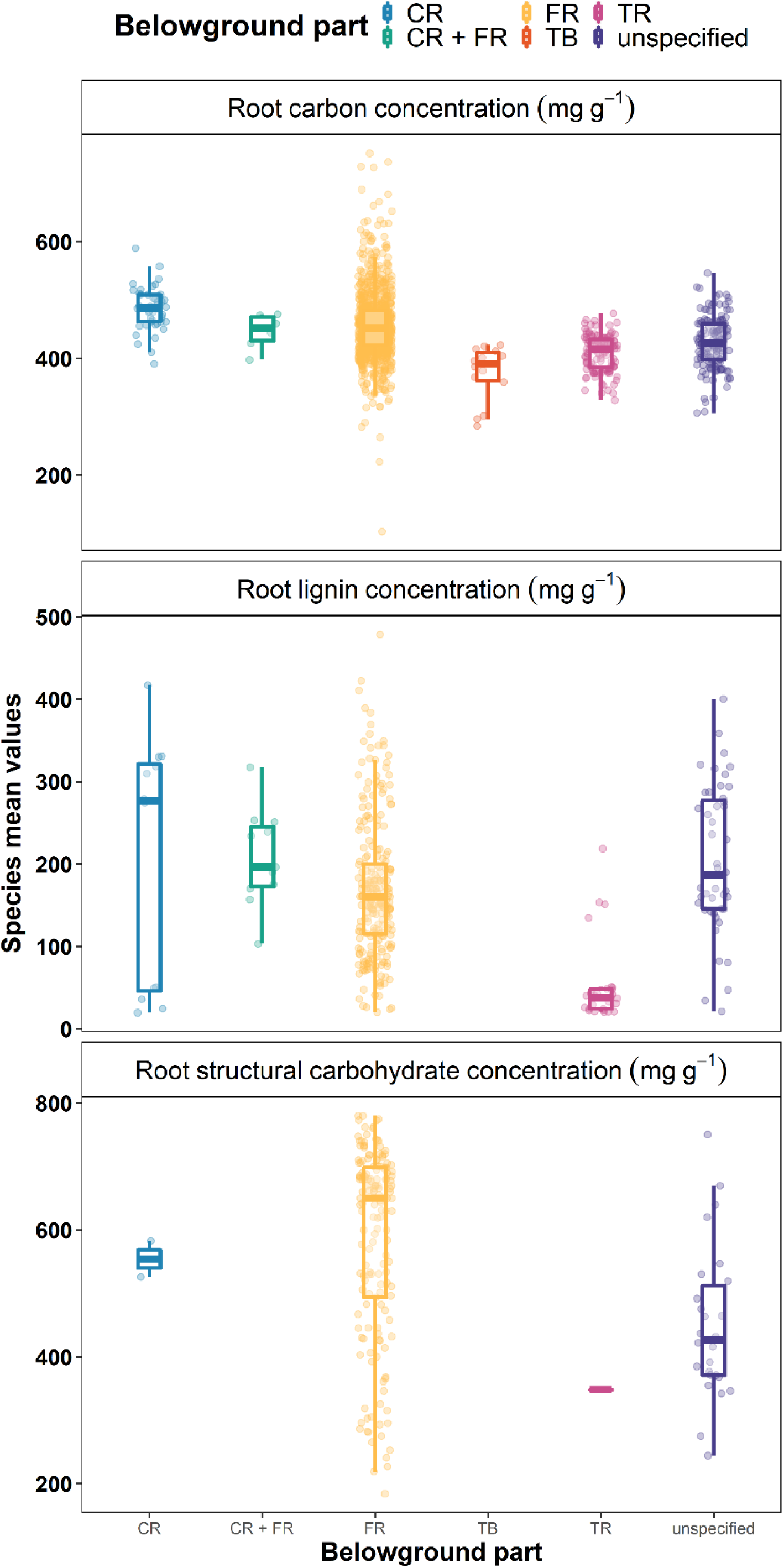
Density plots for chemical traits. Points represent species mean values. Data from coarse (CR), fine (FR) and total roots (TR), total belowground (TB) or unspecified.

**Figure S8.**
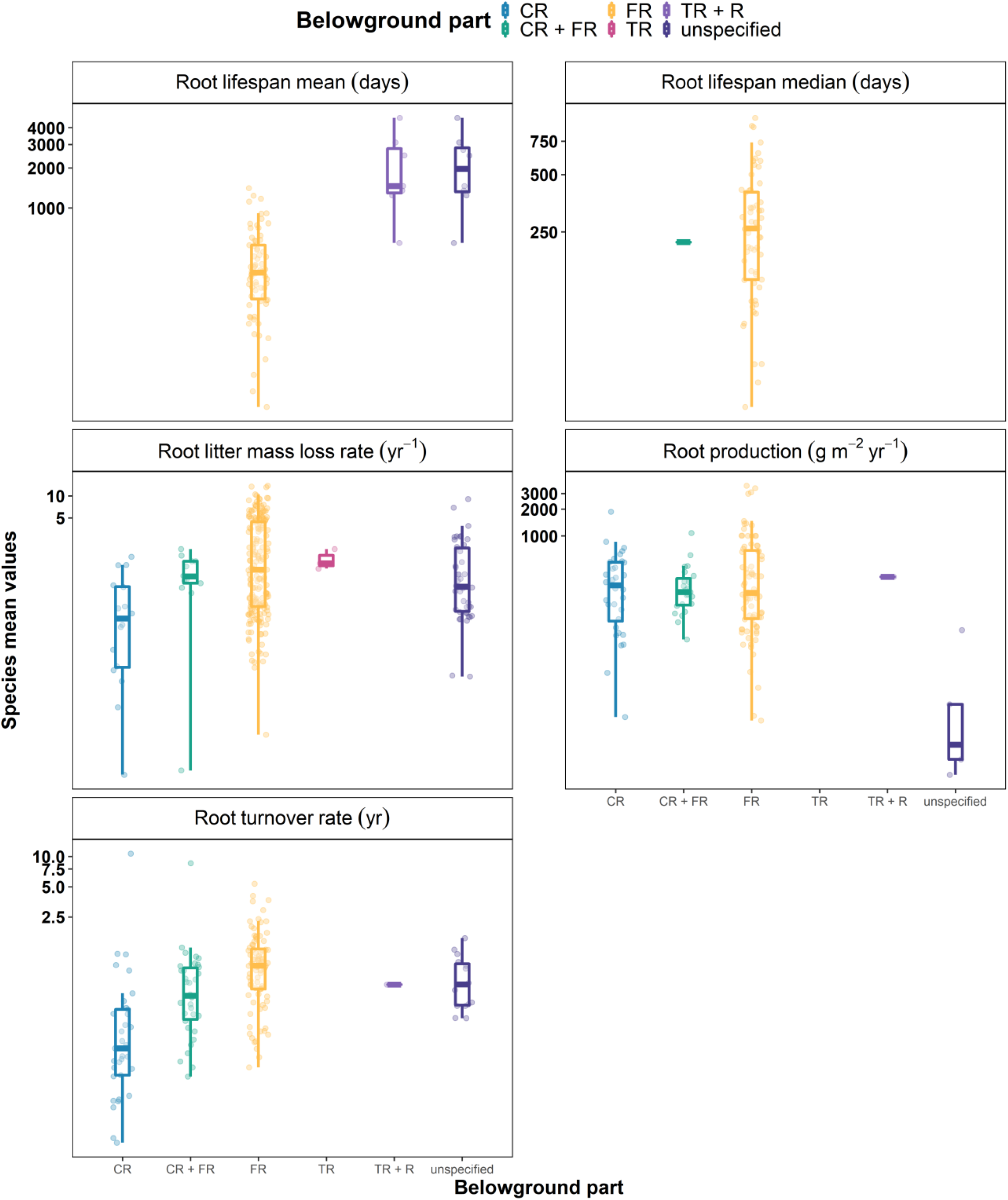
Density plots for dynamics and decomposition. Points represent species mean values. Axis “y” is logarithmic base 2. Data from coarse (CR), fine (FR) and total roots (TR), rhizomes (R), or unspecified.

**Figure S9.**
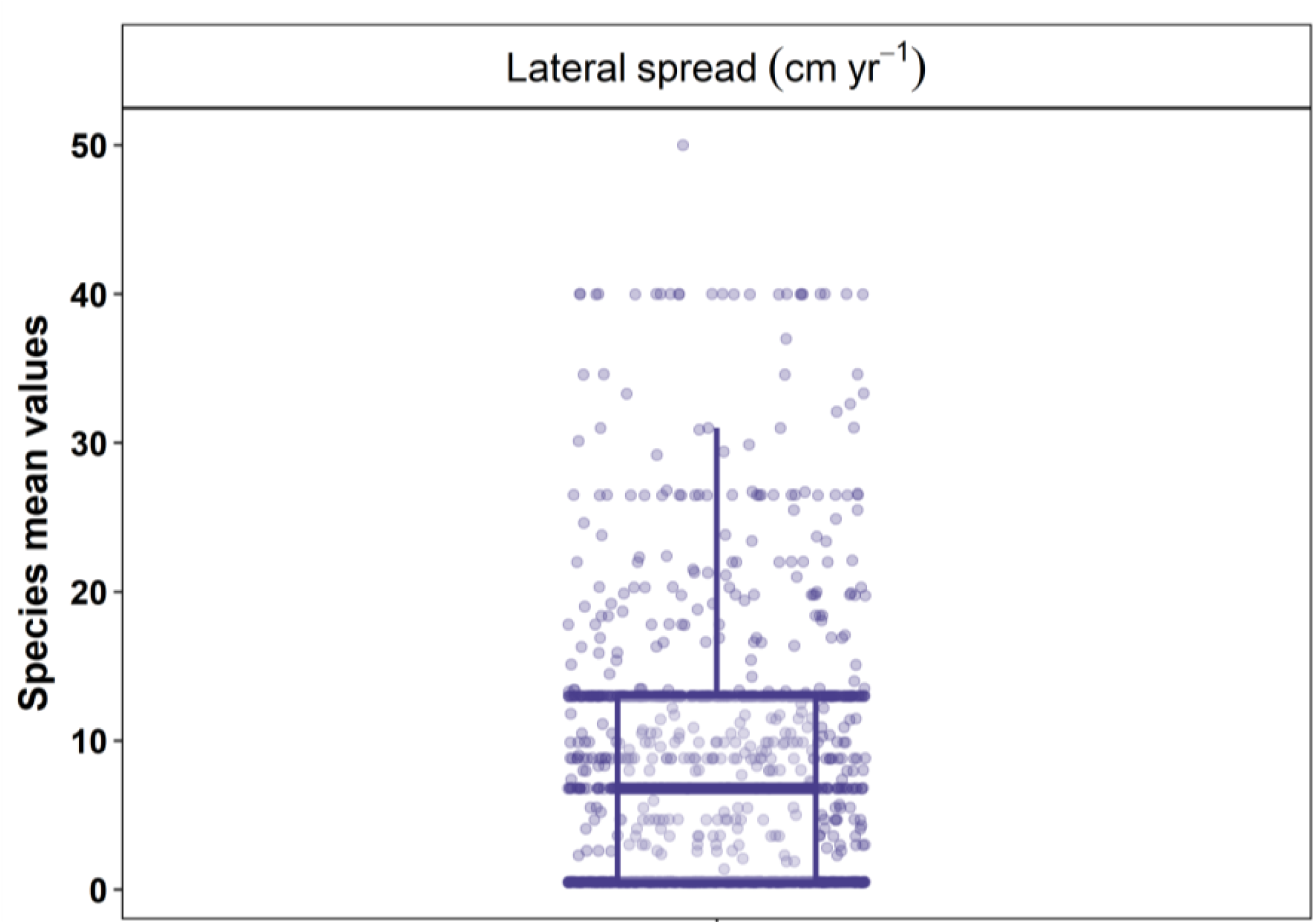
Density plots for lateral spread. Points represent species mean values. Distances are estimated within categories, with mean values of their ranges.

**Figure S10.**
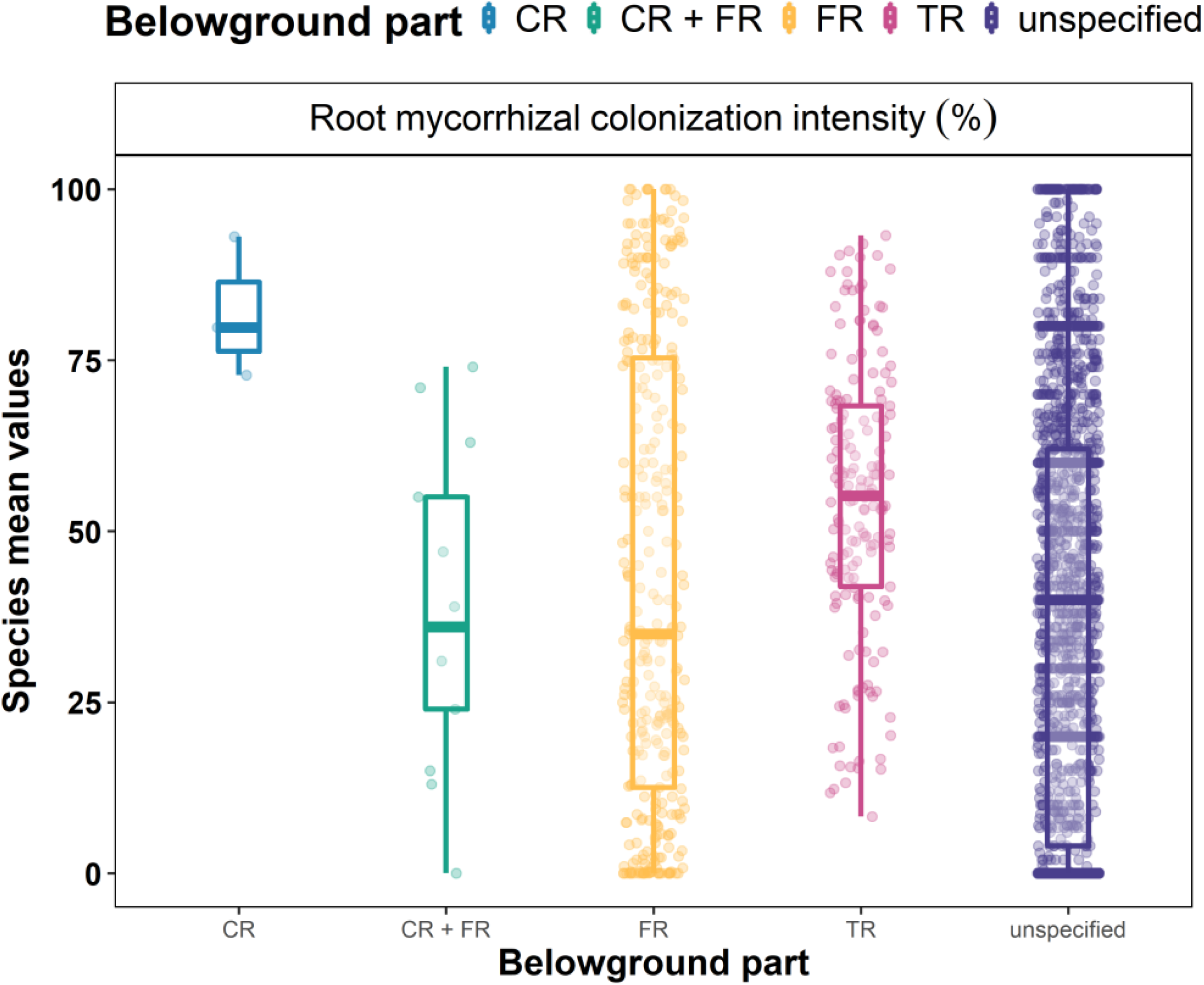
Density plots for root mycorrhizal colonization intensity. Points represent species mean values. Data from coarse (CR), fine (FR) and total roots (TR), or unspecified.

**Figure S11.**
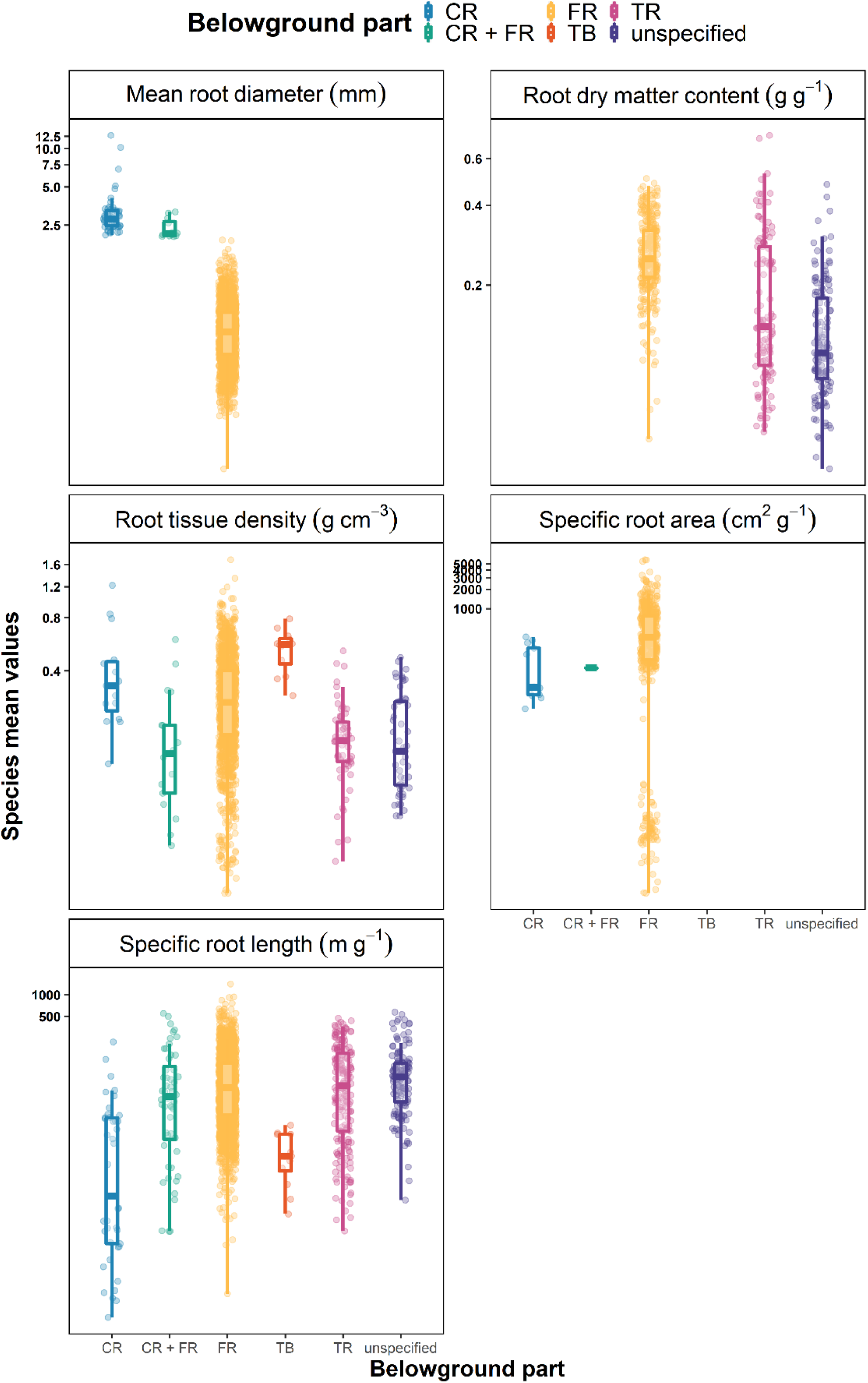
Density plots for morphological traits. Points represent species mean values. Axis “y” is logarithmic base 2. Data from coarse (CR), fine (FR) and total roots (TR), total belowground (TB) or unspecified.

**Figure S12.**
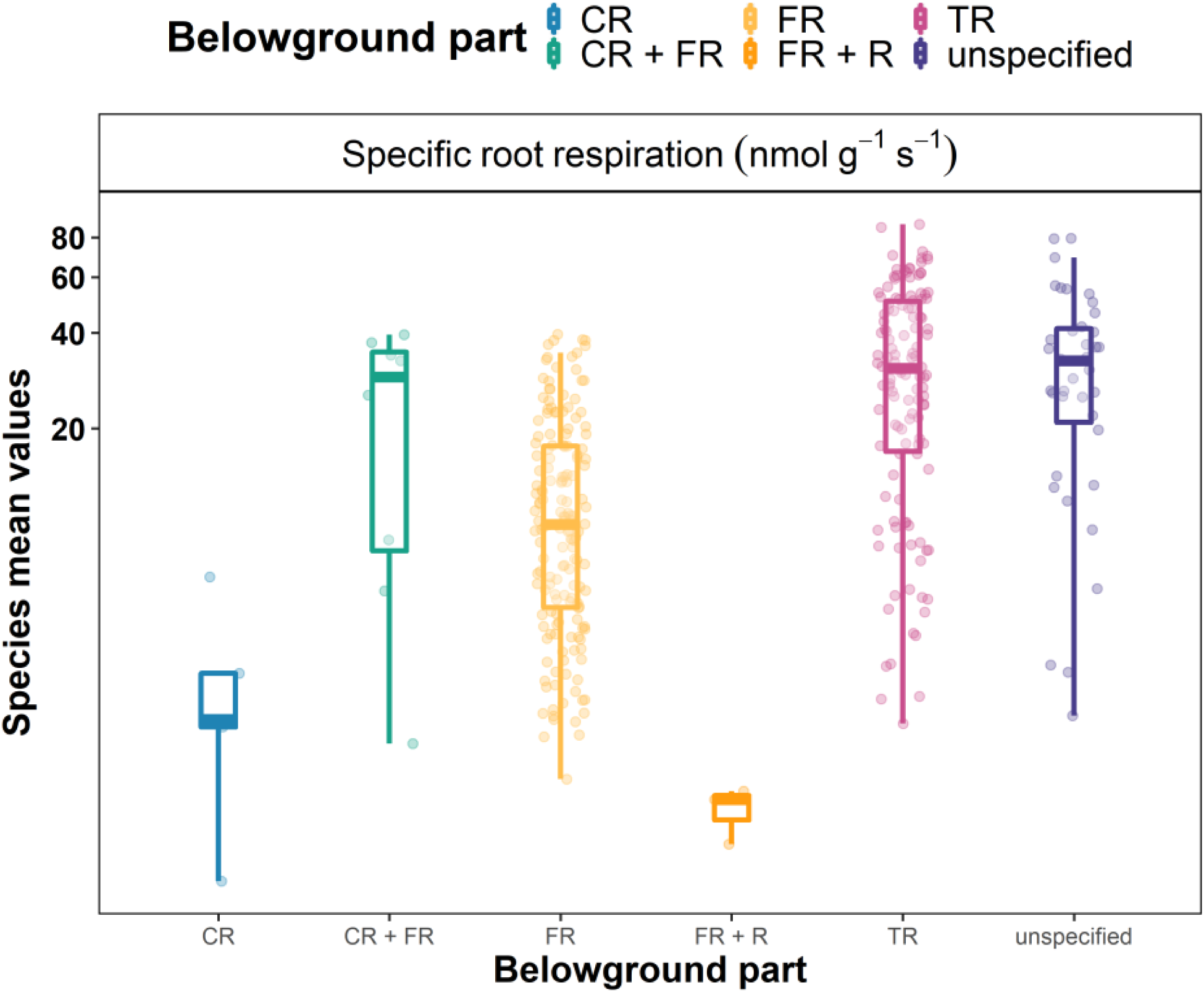
Density plots for specific root respiration. Points represent species mean values. Data from coarse (CR), fine (FR) and total roots (TR), rhizomes (R) or unspecified.

**Figure S13.**
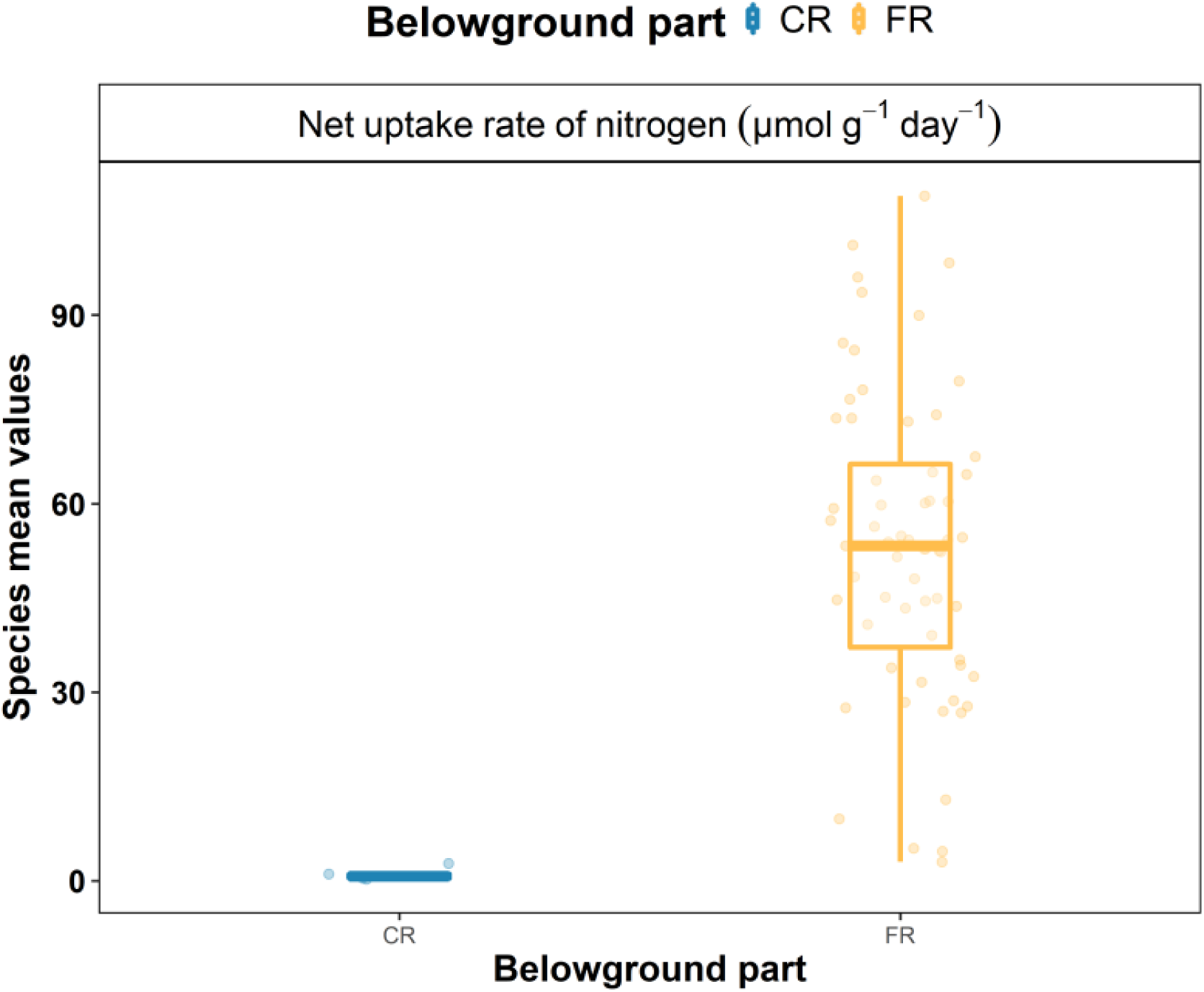
Density plots for net uptake rate of nitrogen. Points represent species mean values. Data from coarse (CR) or fine roots (FR).

**Figure S14.**
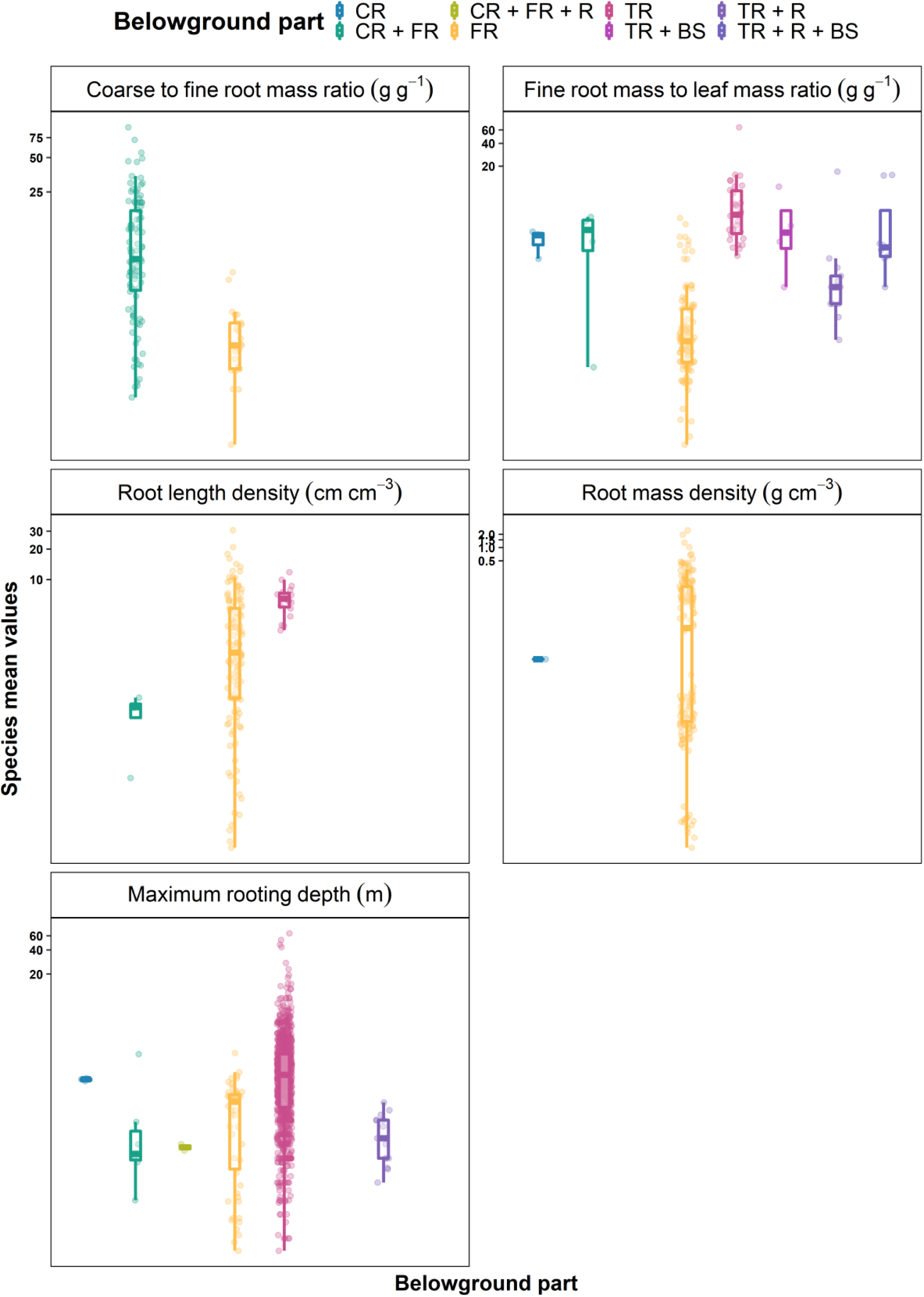
Density plots for system and distribution traits. Points represent species mean values. Axis “y” is logarithmic base 2. Data from coarse (CR), fine (FR) and total roots (TR), belowground steam (BS), and rhizomes (R).

## References

Addo-Danso, S.D., Defrenne, C.E., McCormack, M.L., Ostonen, I., Addo-Danso, A., Foli, E.G., Borden, K.A., Isaac, M.E. & Prescott, C.E. (2019) Fine-root morphological trait variation in tropical forest ecosystems: an evidence synthesis. Plant Ecology.

Adler, P.B., Salguero-Gómez, R., Compagnoni, A., Hsu, J.S., Ray-Mukherjee, J., Mbeau-Ache, C. & Franco, M. (2014) Functional traits explain variation in plant life history strategies. Proceedings of the National Academy of Sciences, 111, 740–745.

Aubin, I., Munson, A.D., Cardou, F., Burton, P.J., Isabel, N., Pedlar, J.H., Paquette, A., Taylor, A.R., Delagrange, S., Kebli, H., Messier, C., Shipley, B., Valladares, F., Kattge, J., Boisvert-Marsh, L. & McKenney, D. (2016) Traits to stay, traits to move: a review of functional traits to assess sensitivity and adaptive capacity of temperate and boreal trees to climate change. Environmental Reviews, 24, 164–186.

Averill, C., Bhatnagar, J.M., Dietze, M.C., Pearse, W.D. & Kivlin, S.N. (2019) Global imprint of mycorrhizal fungi on whole-plant nutrient economics. Proceedings of the National Academy of Sciences, 201906655.

Bardgett, R.D., Mommer, L. & De Vries, F.T. (2014) Going underground: root traits as drivers of ecosystem processes. Trends in Ecology & Evolution, 29, 692–699.

Bergmann, J., Weigelt, A., Plas, F. van der, Laughlin, D.C., Kuyper, T.W., Guerrero-Ramirez, N., Valverde-Barrantes, O.J., Bruelheide, H., Freschet, G.T., Iversen, C.M., Kattge, J., McCormack, M.L., Meier, I.C., Rillig, M.C., Roumet, C., Semchenko, M., Sweeney, C.J., Ruijven, J. van, York, L.M. & Mommer, L. (2020) The fungal collaboration gradient dominates the root economics space in plants. bioRxiv, 2020.01.17.908905.

Boyle, B., Hopkins, N., Lu, Z., Raygoza Garay, J.A., Mozzherin, D., Rees, T., Matasci, N., Narro, M.L., Piel, W.H., McKay, S.J., Lowry, S., Freeland, C., Peet, R.K. & Enquist, B.J. (2013) The taxonomic name resolution service: an online tool for automated standardization of plant names. BMC Bioinformatics, 14, 16.

Breitschwerdt, E., Jandt, U. & Bruelheide, H. (2018) Using co-occurrence information and trait composition to understand individual plant performance in grassland communities. Scientific Reports, 8, 1–15.

Bruelheide, H., Dengler, J., Purschke, O., Lenoir, J., Jiménez-Alfaro, B., Hennekens, S.M., Botta-Dukát, Z., Chytrý, M., Field, R., Jansen, F., Kattge, J., Pillar, V.D., Schrodt, F., Mahecha, M.D., Peet, R.K., Sandel, B., Bodegom, P. van, Altman, J., Alvarez-Dávila, E., Khan, M.A.S.A., Attorre, F., Aubin, I., Baraloto, C., Barroso, J.G., Bauters, M., Bergmeier, E., Biurrun, I., Bjorkman, A.D., Blonder, B., Čarni, A., Cayuela, L., Černý, T., Cornelissen, J.H.C., Craven, D., Dainese, M., Derroire, G., Sanctis, M.D., Díaz, S., Doležal, J., Farfan-Rios, W., Feldpausch, T.R., Fenton, N.J., Garnier, E., Guerin, G.R., Gutiérrez, A.G., Haider, S., Hattab, T., Henry, G., Hérault, B., Higuchi, P., Hölzel, N., Homeier, J., Jentsch, A., Jürgens, N., Kącki, Z., Karger, D.N., Kessler, M., Kleyer, M., Knollová, I., Korolyuk, A.Y., Kühn, I., Laughlin, D.C., Lens, F., Loos, J., Louault, F., Lyubenova, M.I., Malhi, Y., Marcenò, C., Mencuccini, M., Müller, J.V., Munzinger, J., Myers-Smith, I.H., Neill, D.A., Niinemets, Ü., Orwin, K.H., Ozinga, W.A., Penuelas, J., Pérez-Haase, A., Petřík, P., Phillips, O.L., Pärtel, M., Reich, P.B., Römermann, C., Rodrigues, A.V., Sabatini, F.M., Sardans, J., Schmidt, M., Seidler, G., Espejo, J.E.S., Silveira, M., Smyth, A., Sporbert, M., Svenning, J.-C., Tang, Z., Thomas, R., Tsiripidis, I., Vassilev, K., Violle, C., Virtanen, R., Weiher, E., Welk, E., Wesche, K., Winter, M., Wirth, C. & Jandt, U. (2018) Global trait–environment relationships of plant communities. Nature Ecology & Evolution, 2, 1906.

Bryant, C., Wheeler, N.R., Rubel, F. & French, R.H. (2017) kgc: Koeppen-Geiger Climatic Zones, R package version 1.0.0.2,.

Calow, P. (1987) Towards a Definition of Functional Ecology. Functional Ecology, 1, 57–61.

Chamberlain, S.A. & Szöcs, E. (2013) taxize: taxonomic search and retrieval in R. F1000Research, 2.

Chave, J., Coomes, D., Jansen, S., Lewis, S.L., Swenson, N.G. & Zanne, A.E. (2009) Towards a worldwide wood economics spectrum. Ecology Letters, 12, 351–366.

Cornwell, W.K., Pearse, W.D., Dalrymple, R.L. & Zanne, A.E. (2019) What we (don’t) know about global plant diversity. Ecography, 42, 1819–1831.

Craven, D., Eisenhauer, N., Pearse, W.D., Hautier, Y., Isbell, F., Roscher, C., Bahn, M., Beierkuhnlein, C., Bönisch, G., Buchmann, N., Byun, C., Catford, J.A., Cerabolini, B.E.L., Cornelissen, J.H.C., Craine, J.M., Luca, E.D., Ebeling, A., Griffin, J.N., Hector, A., Hines, J., Jentsch, A., Kattge, J., Kreyling, J., Lanta, V., Lemoine, N., Meyer, S.T., Minden, V., Onipchenko, V., Polley, H.W., Reich, P.B., Ruijven, J. van, Schamp, B., Smith, M.D., Soudzilovskaia, N.A., Tilman, D., Weigelt, A., Wilsey, B. & Manning, P. (2018) Multiple facets of biodiversity drive the diversity–stability relationship. Nature Ecology & Evolution, 2, 1579.

Delory, B.M., Weidlich, E.W.A., Meder, L., Lütje, A., Duijnen, R. van, Weidlich, R. & Temperton, V.M. (2017) Accuracy and bias of methods used for root length measurements in functional root research. Methods in Ecology and Evolution, 8, 1594–1606.

Díaz, S. & Cabido, M. (2001) Vive la différence: plant functional diversity matters to ecosystem processes. Trend in Ecology & Evolution, 16, 646–655.

Díaz, S., Kattge, J., Cornelissen, J.H.C., Wright, I.J., Lavorel, S., Dray, S., Reu, B., Kleyer, M., Wirth, C., Colin Prentice, I., Garnier, E., Bönisch, G., Westoby, M., Poorter, H., Reich, P.B., Moles, A.T., Dickie, J., Gillison, A.N., Zanne, A.E., Chave, J., Joseph Wright, S., Sheremet’ev, S.N., Jactel, H., Baraloto, C., Cerabolini, B., Pierce, S., Shipley, B., Kirkup, D., Casanoves, F., Joswig, J.S., Günther, A., Falczuk, V., Rüger, N., Mahecha, M.D. & Gorné, L.D. (2016) The global spectrum of plant form and function. Nature, 529, 167–171.

Freschet, G.T., Cornelissen, J.H.C., van Logtestijn, R.S.P. & Aerts, R. (2010) Evidence of the “plant economics spectrum” in a subartic flora. Journal of Ecology, 98, 362–373.

Freschet, G.T., Pagès, L., Iversen, C.M., Comas, L.H., Rewald, B., Roumet, C., Klimešová, J., Zadworny, M., Poorter, H., Postma, J.A., Adams, T.S., Bagniewska-Zadworna, A., Blancaflor, E.B., Brunner, I., Cornelissen, J.H.C., Garnier, E., Gessler, A., Hobbie, S.E., Lambers, H., Meier, I.C., Mommer, L., Picon-Cochard, C., Rose, L., Ryser, P., Scherer-Lorenzen, M., Soudzilovskaia, N.A., Stokes, A., Sun, T., Valverde-Barrantes, O.J., Weemstra, M., Weigelt, A., Wurzburger, N., York, L.M., Batterman, S.A., Bengough, A.G., Gomes de Moraes, M., Janeček, Š., Salmon, V., Tharayil, N. & McCormack, M.L. (submitted) A starting guide to root ecology: strengthening ecological concepts and standardizing root classification, sampling, processing and trait measurements. New Phytologist.

Freschet, G.T. & Roumet, C. (2017) Sampling roots to capture plant and soil functions. Functional Ecology, 31, 1506–1518.

Freschet, G.T., Valverde-Barrantes, O.J., Tucker, C.M., Craine, J.M., McCormack, M.L., Violle, C., Fort, F., Blackwood, C.B., Urban-Mead, K.R., Iversen, C.M., Bonis, A., Comas, L.H., Cornelissen, J.H.C., Dong, M., Guo, D., Hobbie, S.E., Holdaway, R.J., Kembel, S.W., Makita, N., Onipchenko, V.G., Pico-Cochard, C., Reich, P.B., de la Riva, E.G., Smith, S.W., Soudzilovskaia, N.A., Tjoelker, M.G., Wardle, D.A. & Roumet, C. (2017) Climate, soil and plant functional types as drivers of global fine-root trait variation. Journal of Ecology, 105, 1182–1196.

Gallagher, R.V., Falster, D.S., Maitner, B.S., Salguero-Gómez, R., Vandvik, V., Pearse, W.D., Schneider, F.D., Kattge, J., Alroy, J., Ankenbrand, M.J., Andrew, S.C., Balk, M., Bland, L.M., Boyle, B.L., Bravo-Avila, C.H., Brennan, I., Carthey, A.J.R., Catullo, R., Cavazos, B.R., Chown, S., Fadrique, B., Feng, X., Gibb, H., Halbritter, A.H., Hammock, J., Hogan, J.A., Holewa, H., Hope, M., Iversen, C.M., Jochum, M., Kearney, M., Keller, A., Mabee, P., Madin, J.S., Manning, P., McCormack, L., Michaletz, S.T., Park, D.S., Penone, C., Perez, T.M., Pineda-Munoz, S., Poelen, J., Ray, C.A., Rossetto, M., Sauquet, H., Sparrow, B., Spasojevic, M.J., Telford, R.J., Tobias, J.A., Violle, C., Walls, R., Weiss, K.C.B., Westoby, M., Wright, I.J. & Enquist, B.J. (2019) The Open Traits Network: Using Open Science principles to accelerate trait - based science across the Tree of Life. EcoEvoRvix.

Geber, M.A. & Griffen, L.R. (2003) Inheritance and Natural Selection on Functional Traits. International Journal of Plant Sciences, 164, S21–S42.

Grime, J.P. (1977) Evidence for the existence of three primary strategies in plants and its relevance to ecological and evolutionary theory. The American Naturalist, 111, 1169–1194.

Grime, J.P. (1974) Vegetation classification by reference to strategies. Nature, 250, 26–31.

Iversen, C., Powell, A., McCormack, M., Blackwood, C., Freschet, G., Kattge, J., Roumet, C., Stover, D., Soudzilovskaia, N., Valverde-Barrantes, O., Van Bodegom, P. & Violle, C. (2018) Fine-Root Ecology Database (FRED): A global collection of root trait data with coincident site, vegetation, edaphic, and climatic data, Version 2, Oak Ridge National Laboratory, TES SFA, U.S. Department of Energy, Oak Ridge, Tennessee, U.S.A.

Iversen, C.M., McCormack, M.L., Powell, A.S., Blackwood, C.B., Freschet, G.T., Kattge, J., Roumet, C., Stover, D.B., Soudzilovskaia, N.A., Valverde-Barrantes, O.J., van Bodegom, P.M. & Violle, C. (2017) A global Fine-Root Ecology Database to address below-ground challenges in plant ecology. New Phytologist, 215, 15–26.

Kattge, J., Bönisch, G., Díaz, S., Lavorel, S., Prentice, I.C., Leadley, P., Tautenhahn, S., Werner, G.D.A., Aakala, T., Abedi, M., Acosta, A.T.R., Adamidis, G.C., Adamson, K., Aiba, M., Albert, C.H., Alcántara, J.M., c, C.A., Aleixo, I., Ali, H., Amiaud, B., Ammer, C., Amoroso, M.M., Anand, M., Anderson, C., Anten, N., Antos, J., Apgaua, D.M.G., Ashman, T.-L., Asmara, D.H., Asner, G.P., Aspinwall, M., Atkin, O., Aubin, I., Baastrup-Spohr, L., Bahalkeh, K., Bahn, M., Baker, T., Baker, W.J., Bakker, J.P., Baldocchi, D., Baltzer, J., Banerjee, A., Baranger, A., Barlow, J., Barneche, D.R., Baruch, Z., Bastianelli, D., Battles, J., Bauerle, W., Bauters, M., Bazzato, E., Beckmann, M., Beeckman, H., Beierkuhnlein, C., Bekker, R., Belfry, G., Belluau, M., Beloiu, M., Benavides, R., Benomar, L., Berdugo-Lattke, M.L., Berenguer, E., Bergamin, R., Bergmann, J., Carlucci, M.B., Berner, L., Bernhardt-Römermann, M., Bigler, C., Bjorkman, A.D., Blackman, C., Blanco, C., Blonder, B., Blumenthal, D., Bocanegra-González, K.T., Boeckx, P., Bohlman, S., Böhning-Gaese, K., Boisvert-Marsh, L., Bond, W., Bond-Lamberty, B., Boom, A., Boonman, C.C.F., Bordin, K., Boughton, E.H., Boukili, V., Bowman, D.M.J.S., Bravo, S., Brendel, M.R., Broadley, M.R., Brown, K.A., Bruelheide, H., Brumnich, F., Bruun, H.H., Bruy, D., Buchanan, S.W., Bucher, S.F., Buchmann, N., Buitenwerf, R., Bunker, D.E., Bürger, J., Burrascano, S., Burslem, D.F.R.P., Butterfield, B.J., Byun, C., Marques, M., Scalon, M.C., Caccianiga, M., Cadotte, M., Cailleret, M., Camac, J., Camarero, J.J., Campany, C., Campetella, G., Campos, J.A., Cano-Arboleda, L., Canullo, R., Carbognani, M., Carvalho, F., Casanoves, F., Castagneyrol, B., Catford, J.A., Cavender-Bares, J., Cerabolini, B.E.L., Cervellini, M., Chacón-Madrigal, E., Chapin, K., Chapin, F.S., Chelli, S., Chen, S.-C., Chen, A., Cherubini, P., Chianucci, F., Choat, B., Chung, K.-S., Chytrý, M., Ciccarelli, D., Coll, L., Collins, C.G., Conti, L., Coomes, D., Cornelissen, J.H.C., Cornwell, W.K., Corona, P., Coyea, M., Craine, J., Craven, D., Cromsigt, J.P.G.M., Csecserits, A., Cufar, K., Cuntz, M., Silva, A.C. da, Dahlin, K.M., Dainese, M., Dalke, I., Fratte, M.D., Dang-Le, A.T., Danihelka, J., Dannoura, M., Dawson, S., Beer, A.J. de, Frutos, A.D., Long, J.R.D., Dechant, B., Delagrange, S., Delpierre, N., Derroire, G., Dias, A.S., Diaz-Toribio, M.H., Dimitrakopoulos, P.G., Dobrowolski, M., Doktor, D., Dřevojan, P., Dong, N., Dransfield, J., Dressler, S., Duarte, L., Ducouret, E., Dullinger, S., Durka, W., Duursma, R., Dymova, O., E-Vojtkó, A., Eckstein, R.L., Ejtehadi, H., Elser, J., Emilio, T., Engemann, K., Erfanian, M.B., Erfmeier, A., Esquivel-Muelbert, A., Esser, G., Estiarte, M., Domingues, T.F., Fagan, W.F., Fagúndez, J., Falster, D.S., Fan, Y., Fang, J., Farris, E., Fazlioglu, F., Feng, Y., Fernandez-Mendez, F., Ferrara, C., Ferreira, J., Fidelis, A., Finegan, B., Firn, J., Flowers, T.J., Flynn, D.F.B., Fontana, V., Forey, E., Forgiarini, C., François, L., Frangipani, M., Frank, D., Frenette-Dussault, C., Freschet, G.T., Fry, E.L., Fyllas, N.M., Mazzochini, G.G., Gachet, S., Gallagher, R., Ganade, G., Ganga, F., García-Palacios, P., Gargaglione, V., Garnier, E., Garrido, J.L., Gasper, A.L. de, Gea-Izquierdo, G., Gibson, D., Gillison, A.N., Giroldo, A., Glasenhardt, M.-C., Gleason, S., Gliesch, M., Goldberg, E., Göldel, B., Gonzalez-Akre, E., Gonzalez-Andujar, J.L., González-Melo, A., González-Robles, A., Graae, B.J., Granda, E., Graves, S., Green, W.A., Gregor, T., Gross, N., Guerin, G.R., Günther, A., Gutiérrez, A.G., Haddock, L., Haines, A., Hall, J., Hambuckers, A., Han, W., Harrison, S.P., Hattingh, W., Hawes, J.E., He, T., He, P., Heberling, J.M., Helm, A., Hempel, S., Hentschel, J., Hérault, B., Hereş, A.-M., Herz, K., Heuertz, M., Hickler, T., Hietz, P., Higuchi, P., Hipp, A.L., Hirons, A., Hock, M., Hogan, J.A., Holl, K., Honnay, O., Hornstein, D., Hou, E., Hough-Snee, N., Hovstad, K.A., Ichie, T., Igić, B., Illa, E., Isaac, M., Ishihara, M., Ivanov, L., Ivanova, L., Iversen, C.M., Izquierdo, J., Jackson, R.B., Jackson, B., Jactel, H., Jagodzinski, A.M., Jandt, U., Jansen, S., Jenkins, T., Jentsch, A., Jespersen, J.R.P., Jiang, G.-F., Johansen, J.L., Johnson, D., Jokela, E.J., Joly, C.A., Jordan, G.J., Joseph, G.S., Junaedi, D., Junker, R.R., Justes, E., Kabzems, R., Kane, J., Kaplan, Z., Kattenborn, T., Kavelenova, L., Kearsley, E., Kempel, A., Kenzo, T., Kerkhoff, A., Khalil, M.I., Kinlock, N.L., Kissling, W.D., Kitajima, K., Kitzberger, T., Kjøller, R., Klein, T., Kleyer, M., Klimešová, J., Klipel, J., Kloeppel, B., Klotz, S., Knops, J.M.H., Kohyama, T., Koike, F., Kollmann, J., Komac, B., Komatsu, K., König, C., Kraft, N.J.B., Kramer, K., Kreft, H., Kühn, I., Kumarathunge, D., Kuppler, J., Kurokawa, H., Kurosawa, Y., Kuyah, S., Laclau, J.-P., Lafleur, B., Lallai, E., Lamb, E., Lamprecht, A., Larkin, D.J., Laughlin, D., Bagousse-Pinguet, Y.L., Maire, G. le, Roux, P.C. le, Roux, E. le, Lee, T., Lens, F., Lewis, S.L., Lhotsky, B., Li, Y., Li, X., Lichstein, J.W., Liebergesell, M., Lim, J.Y., Lin, Y.-S., Linares, J.C., Liu, C., Liu, D., Liu, U., Livingstone, S., Llusià, J., Lohbeck, M., López-García, Á., Lopez-Gonzalez, G., Lososová, Z., Louault, F., Lukács, B.A., Lukeš, P., Luo, Y., Lussu, M., Ma, S., Pereira, C.M.R., Mack, M., Maire, V., Mäkelä, A., Mäkinen, H., Malhado, A.C.M., Mallik, A., Manning, P., Manzoni, S., Marchetti, Z., Marchino, L., Marcilio-Silva, V., Marcon, E., Marignani, M., Markesteijn, L., Martin, A., Martínez-Garza, C., Martínez-Vilalta, J., Mašková, T., Mason, K., Mason, N., Massad, T.J., Masse, J., Mayrose, I., McCarthy, J., McCormack, M.L., McCulloh, K., McFadden, I.R., McGill, B.J., McPartland, M.Y., Medeiros, J.S., Medlyn, B., Meerts, P., Mehrabi, Z., Meir, P., Melo, F.P.L., Mencuccini, M., Meredieu, C., Messier, J., Mészáros, I., Metsaranta, J., Michaletz, S.T., Michelaki, C., Migalina, S., Milla, R., Miller, J.E.D., Minden, V., Ming, R., Mokany, K., Moles, A.T., Molnár, A., Molofsky, J., Molz, M., Montgomery, R.A., Monty, A., Moravcová, L., Moreno-Martínez, A., Moretti, M., Mori, A.S., Mori, S., Morris, D., Morrison, J., Mucina, L., Mueller, S., Muir, C.D., Müller, S.C., Munoz, F., Myers-Smith, I.H., Myster, R.W., Nagano, M., Naidu, S., Narayanan, A., Natesan, B., Negoita, L., Nelson, A.S., Neuschulz, E.L., Ni, J., Niedrist, G., Nieto, J., Niinemets, Ü., Nolan, R., Nottebrock, H., Nouvellon, Y., Novakovskiy, A., Nystuen, K.O., O’Grady, A., O’Hara, K., O’Reilly-Nugent, A., Oakley, S., Oberhuber, W., Ohtsuka, T., Oliveira, R., Öllerer, K., Olson, M.E., Onipchenko, V., Onoda, Y., Onstein, R.E., Ordonez, J.C., Osada, N., Ostonen, I., Ottaviani, G., Otto, S., Overbeck, G.E., Ozinga, W.A., Pahl, A.T., Paine, C.E.T., Pakeman, R.J., Papageorgiou, A.C., Parfionova, E., Pärtel, M., Patacca, M., Paula, S., Paule, J., Pauli, H., Pausas, J.G., Peco, B., Penuelas, J., Perea, A., Peri, P.L., Petisco-Souza, A.C., Petraglia, A., Petritan, A.M., Phillips, O.L., Pierce, S., Pillar, V.D., Pisek, J., Pomogaybin, A., Poorter, H., Portsmuth, A., Poschlod, P., Potvin, C., Pounds, D., Powell, A.S., Power, S.A., Prinzing, A., Puglielli, G., Pyšek, P., Raevel, V., Rammig, A., Ransijn, J., Ray, C.A., Reich, P.B., Reichstein, M., Reid, D.E.B., Réjou-Méchain, M., Dios, V.R. de, Ribeiro, S., Richardson, S., Riibak, K., Rillig, M.C., Riviera, F., Robert, E.M.R., Roberts, S., Robroek, B., Roddy, A., Rodrigues, A.V., Rogers, A., Rollinson, E., Rolo, V., Römermann, C., Ronzhina, D., Roscher, C., Rosell, J.A., Rosenfield, M.F., Rossi, C., Roy, D.B., Royer-Tardif, S., Rüger, N., Ruiz-Peinado, R., Rumpf, S.B., Rusch, G.M., Ryo, M., Sack, L., Saldaña, A., Salgado-Negret, B., Salguero-Gomez, R., Santa-Regina, I., Santacruz-García, A.C., Santos, J., Sardans, J., Schamp, B., Scherer-Lorenzen, M., Schleuning, M., Schmid, B., Schmidt, M., Schmitt, S., Schneider, J.V., Schowanek, S.D., Schrader, J., Schrodt, F., Schuldt, B., Schurr, F., Garvizu, G.S., Semchenko, M., Seymour, C., Sfair, J.C., Sharpe, J.M., Sheppard, C.S., Sheremetiev, S., Shiodera, S., Shipley, B., Shovon, T.A., Siebenkäs, A., Sierra, C., Silva, V., Silva, M., Sitzia, T., Sjöman, H., Slot, M., Smith, N.G., Sodhi, D., Soltis, P., Soltis, D., Somers, B., Sonnier, G., Sørensen, M.V., Sosinski, E.E., Soudzilovskaia, N.A., Souza, A.F., Spasojevic, M., Sperandii, M.G., Stan, A.B., Stegen, J., Steinbauer, K., Stephan, J.G., Sterck, F., Stojanovic, D.B., Strydom, T., Suarez, M.L., Svenning, J.-C., Svitková, I., Svitok, M., Svoboda, M., Swaine, E., Swenson, N., Tabarelli, M., Takagi, K., Tappeiner, U., Tarifa, R., Tauugourdeau, S., Tavsanoglu, C., Beest, M. te, Tedersoo, L., Thiffault, N., Thom, D., Thomas, E., Thompson, K., Thornton, P.E., Thuiller, W., Tichý, L., Tissue, D., Tjoelker, M.G., Tng, D.Y.P., Tobias, J., Török, P., Tarin, T., Torres-Ruiz, J.M., Tóthmérész, B., Treurnicht, M., Trivellone, V., Trolliet, F., Trotsiuk, V., Tsakalos, J.L., Tsiripidis, I., Tysklind, N., Umehara, T., Usoltsev, V., Vadeboncoeur, M., Vaezi, J., Valladares, F., Vamosi, J., Bodegom, P.M. van, Breugel, M. van, Cleemput, E.V., Weg, M. van de, Merwe, S. van der, Plas, F. van der, Sande, M.T. van der, Kleunen, M. van, Meerbeek, K.V., Vanderwel, M., Vanselow, K.A., Vårhammar, A., Varone, L., Valderrama, M.Y.V., Vassilev, K., Vellend, M., Veneklaas, E.J., Verbeeck, H., Verheyen, K., Vibrans, A., Vieira, I., Villacís, J., Violle, C., Vivek, P., Wagner, K., Waldram, M., Waldron, A., Walker, A.P., Waller, M., Walther, G., Wang, H., Wang, F., Wang, W., Watkins, H., Watkins, J., Weber, U., Weedon, J.T., Wei, L., Weigelt, P., Weiher, E., Wells, A.W., Wellstein, C., Wenk, E., Westoby, M., Westwood, A., White, P.J., Whitten, M., Williams, M., Winkler, D.E., Winter, K., Womack, C., Wright, I.J., Wright, S.J., Wright, J., Pinho, B.X., Ximenes, F., Yamada, T., Yamaji, K., Yanai, R., Yankov, N., Yguel, B., Zanini, K.J., Zanne, A.E., Zelený, D., Zhao, Y.-P., Zheng, J., Zheng, J., Ziemińska, K., Zirbel, C.R., Zizka, G., Zo-Bi, I.C., Zotz, G. & Wirth, C. (2020) TRY plant trait database – enhanced coverage and open access. Global Change Biology, 26, 119–188.

Kattge, J., Díaz, S., Lavorel, S., Prentice, I.C., Leadley, P., Bönisch, G., Garnier, E., Westoby, M., Reich, P.B., Wright, I.J., Cornelissen, J.H.C., Violle, C., Harrison, S.P., Bodegom, P.M.V., Reichstein, M., Enquist, B.J., Soudzilovskaia, N.A., Ackerly, D.D., Anand, M., Atkin, O., Bahn, M., Baker, T.R., Baldocchi, D., Bekker, R., Blanco, C.C., Blonder, B., Bond, W.J., Bradstock, R., Bunker, D.E., Casanoves, F., Cavender-Bares, J., Chambers, J.Q., Iii, F.S.C., Chave, J., Coomes, D., Cornwell, W.K., Craine, J.M., Dobrin, B.H., Duarte, L., Durka, W., Elser, J., Esser, G., Estiarte, M., Fagan, W.F., Fang, J., Fernández-Méndez, F., Fidelis, A., Finegan, B., Flores, O., Ford, H., Frank, D., Freschet, G.T., Fyllas, N.M., Gallagher, R.V., Green, W.A., Gutierrez, A.G., Hickler, T., Higgins, S.I., Hodgson, J.G., Jalili, A., Jansen, S., Joly, C.A., Kerkhoff, A.J., Kirkup, D., Kitajima, K., Kleyer, M., Klotz, S., Knops, J.M.H., Kramer, K., Kühn, I., Kurokawa, H., Laughlin, D., Lee, T.D., Leishman, M., Lens, F., Lenz, T., Lewis, S.L., Lloyd, J., Llusià, J., Louault, F., Ma, S., Mahecha, M.D., Manning, P., Massad, T., Medlyn, B.E., Messier, J., Moles, A.T., Müller, S.C., Nadrowski, K., Naeem, S., Niinemets, Ü., Nöllert, S., Nüske, A., Ogaya, R., Oleksyn, J., Onipchenko, V.G., Onoda, Y., Ordoñez, J., Overbeck, G., Ozinga, W.A., Patiño, S., Paula, S., Pausas, J.G., Peñuelas, J., Phillips, O.L., Pillar, V., Poorter, H., Poorter, L., Poschlod, P., Prinzing, A., Proulx, R., Rammig, A., Reinsch, S., Reu, B., Sack, L., Salgado-Negret, B., Sardans, J., Shiodera, S., Shipley, B., Siefert, A., Sosinski, E., Soussana, J.-F., Swaine, E., Swenson, N., Thompson, K., Thornton, P., Waldram, M., Weiher, E., White, M., White, S., Wright, S.J., Yguel, B., Zaehle, S., Zanne, A.E. & Wirth, C. (2011) TRY – a global database of plant traits. Global Change Biology, 17, 2905–2935.

Keddy, P.A. (1992) Assembly and response rules: two goals for predictive community ecology. Journal of Vegetation Science, 3, 157–164.

Klimešová, J. & Bello, F.D. (2009) CLO-PLA: the database of clonal and bud bank traits of Central European flora§. Journal of Vegetation Science, 20, 511–516.

Klimešová, J., Danihelka, J., Chrtek, J., Bello, F. de & Herben, T. (2017) CLO-PLA: a database of clonal and bud-bank traits of the Central European flora. Ecology, 98, 1179–1179.

Klimešová, J., Martínková, J., Pausas, J.G., de Moraes, M.G., Herben, T., Yu, F.-H., Puntieri, J., Vesk, P.A., de Bello, F., Janeček, Š., Altman, J., Appezzato-da-Glória, B., Bartušková, A., Crivellaro, A., Doležal, J., Ott, J.P., Paula, S., Schnablová, R., Schweingruber, F.H. & Ottaviani, G. (2019) Handbook of standardized protocols for collecting plant modularity traits. Perspectives in Plant Ecology, Evolution and Systematics, 40, 125485.

Kong, D., Wang, J., Wu, H., Valverde-Barrantes, O.J., Wang, R., Zeng, H., Kardol, P., Zhang, H. & Feng, Y. (2019) Nonlinearity of root trait relationships and the root economics spectrum. Nature Communications, 10, 2203.

Laliberté, E. (2016) Below-ground frontiers in trait-based plant ecology. New Phytologist, 213, 1597–1603.

Lavorel, S. & Garnier, E. (2002) Predicting changes in community composition and ecosystem functioning from plant traits: revisiting the Holy Grail. Functional Ecology, 16, 545–556.

Ma, Z., Guo, D., Xu, X., Lu, M., Bardgett, R.D., Eissenstat, D.M., McCormack, M.L. & Hedin, L.O. (2018) Evolutionary history resolves global organization of root functional traits. Nature, 555, 94–97.

Martin, A.R. & Isaac, M.E. (2018) Functional traits in agroecology: Advancing description and prediction in agroecosystems. Journal of Applied Ecology, 55, 5–11.

Martin, L.J., Blossey, B. & Ellis, E. (2012) Mapping where ecologists work: biases in the global distribution of terrestrial ecological observations. Frontiers in Ecology and the Environment, 10, 195–201.

McCormack, M.L., Dickie, I.A., Eissenstat, D.M., Fahey, T.J., Fernandez, C.W., Guo, D., Helmisaari, H.-S., Hobbie, E.A., Iversen, C.M., Jackson, R.B., Leppälammi-Kujansuu, J., Norby, R.J., Phillips, R.P., Pregitzer, K.S., Pritchard, S.G., Rewald, B. & Zadworny, M. (2015) Redefining fine roots improves understanding of below-ground contributions to terrestrial biosphere processes. New Phytologist, 207, 505–518.

Minden, V. & Olde Venterink, H. (2019) Plant traits and species interactions along gradients of N, P and K availabilities. Functional Ecology, 33, 1611–1626.

Pausas, J.G., Bradstock, R.A., Keith, D.A. & Keeley, J.E. (2004) Plant Functional Traits in Relation to Fire in Crown-Fire Ecosystems. Ecology, 85, 1085–1100.

Reich, P.B. (2014) The world-wide ‘fast–slow’ plant economics spectrum: a traits manifesto. Journal of Ecology, 102, 275–301.

Reich, P.B., Wright, I.J., Cavender-Bares, J., Craine, J.M., Oleksyn, J., Westoby, M. & Walters, M.B. (2003) The Evolution of Plant Functional Variation: Traits, Spectra, and Strategies. International Journal of Plant Sciences, 164, S143–S164.

Soudzilovskaia, N.A., Vaessen, S., Barcelo, M., He, J., Rahimlou, S., Abarenkov, K., Brundrett, M.C., Gomes, S.I., Merckx, V. & Tedersoo, L. FungalRoot: Global online database of plant mycorrhizal associations. New Phytologist, n/a.

Tedersoo, L., Laanisto, L., Rahimlou, S., Toussaint, A., Hallikma, T. & Pärtel, M. (2018) Global database of plants with root-symbiotic nitrogen fixation: NodDB. Journal of Vegetation Science, 29, 560–568.

Valverde-Barrantes, O.J., Freschet, G.T., Roumet, C. & Blackwood, C.B. (2017) A worldview of root traits: the influence of ancestry, growth form, climate and mycorrhizal association on the functional trait variation of fine-root tissues in seed plants. New Phytologist, 215, 1562–1573.

Violle, C., Navas, M.-L., Vile, D., Kazakou, E., Fortunel, C., Hummel, I. & Garnier, E. (2007) Let the concept of trait be functional! Oikos, 116, 882–892.

Violle, C., Reich, P.B., Pacala, S.W., Enquist, B.J. & Kattge, J. (2014) The emergence and promise of functional biogeography. Proceedings of the National Academy of Sciences, 111, 13690–13696.

Weigelt, P., König, C. & Kreft, H. (2019) GIFT – A Global Inventory of Floras and Traits for macroecology and biogeography. Journal of Biogeography, 47, 16–43.

Wieczynski, D.J., Boyle, B., Buzzard, V., Duran, S.M., Henderson, A.N., Hulshof, C.M., Kerkhoff, A.J., McCarthy, M.C., Michaletz, S.T., Swenson, N.G., Asner, G.P., Bentley, L.P., Enquist, B.J. & Savage, V.M. (2019) Climate shapes and shifts functional biodiversity in forests worldwide. Proceedings of the National Academy of Sciences, 116, 587–592.

Wood, S.A., Karp, D.S., DeClerck, F., Kremen, C., Naeem, S. & Palm, C.A. (2015) Functional traits in agriculture: agrobiodiversity and ecosystem services. Trends in Ecology & Evolution, 30, 531–539.

Wright, I.J., Reich, P.B., Westoby, M., Ackerly, D.D., Baruch, Z., Bongers, F., Cavender-Bares, J., Chapin, T., Cornelissen, J.H.C., Diemer, M., Flexas, J., Garnier, E., Groom, P.K., Gulias, J., Hikosaka, K., Lamont, B.B., Lee, T., Lee, W., Lusk, C., Midgley, J.J., Navas, M.-L., Niinemets, U., Oleksyn, J., Osada, N., Poorter, H., Poot, P., Prior, L., Pyankov, V.I., Roumet, C., Thomas, S.C., Tjoelker, M.G., Veneklaas, E.J. & Villar, R. (2004) The worldwide leaf economics spectrum. Nature, 428, 821–827.

## References

McCormack, M.L., Guo, D., Iversen, C.M., Chen, W., Eissenstat, D.M., Fernandez, C.W., Li, L., Ma, C., Ma, Z., Poorter, H., Reich, P.B., Zadworny, M. & Zanne, A. (2017) Building a better foundation: improving root-trait measurements to understand and model plant and ecosystem processes. New Phytologist, 215, 27–37.

